# Distribution of ubiquilin 2 and TDP-43 aggregates throughout the CNS in *UBQLN2* p.T487I-linked amyotrophic lateral sclerosis and frontotemporal dementia

**DOI:** 10.1101/2023.02.10.527924

**Authors:** Laura R. Nementzik, Kyrah M. Thumbadoo, Helen C. Murray, David Gordon, Shu Yang, Ian P. Blair, Clinton Turner, Richard L. M. Faull, Maurice A. Curtis, Catriona McLean, Garth A. Nicholson, Molly E. V. Swanson, Emma L. Scotter

**Affiliations:** School of Biological Sciences, University of Auckland, Auckland, New Zealand; Centre for Brain Research, University of Auckland, Auckland, New Zealand; Department of Anatomy and Medical Imaging, University of Auckland, Auckland, New Zealand; Centre for Motor Neuron Disease Research, Department of Biomedical Sciences, Faculty of Medicine, Health, and Human Sciences, Macquarie University, North Ryde, NSW, Australia; Department of Anatomical Pathology, LabPlus, Auckland City Hospital, Auckland, New Zealand; Department of Anatomical Pathology, Alfred Health, Melbourne, VIC, Australia; Northcott Neuroscience Laboratory, ANZAC Research Institute, Sydney, Australia; Molecular Medicine Laboratory, Concord Repatriation General Hospital, Sydney, Australia

**Keywords:** *UBQLN2*, ubiquilin 2, amyotrophic lateral sclerosis (ALS), frontotemporal dementia (FTD), neuropathology, TDP-43

## Abstract

Mutations in the *UBQLN2* gene cause amyotrophic lateral sclerosis (ALS) and frontotemporal dementia (FTD). The neuropathology of such *UBQLN2*-linked cases of ALS/FTD is characterised by aggregates of the ubiquilin 2 protein in addition to aggregates of the transactive response DNA-binding protein of 43 kDa (TDP-43). ALS and FTD without *UBQLN2* mutations are also characterised by TDP-43 aggregates, that may or may not colocalise with wildtype ubiquilin 2. Despite this, the relative contributions of TDP-43 and ubiquilin 2 to disease pathogenesis remain largely under-characterised, as does their relative deposition as aggregates across the central nervous system (CNS). Here we conducted multiplex immunohistochemistry of three *UBQLN2* p.T487I-linked ALS/FTD cases, three non-*UBQLN2*-linked (sporadic) ALS cases, and eight non-neurodegenerative disease controls, covering 40 CNS regions. We then quantified ubiquilin 2 aggregates, TDP-43 aggregates, and aggregates containing both proteins in regions of interest to determine how *UBQLN2*-linked and non-*UBQLN2*-linked proteinopathy differ. We find that ubiquilin 2 aggregates that are negative for TDP-43 are predominantly small and punctate, and are abundant in the hippocampal formation, spinal cord, all tested regions of neocortex, medulla, and substantia nigra in *UBQLN2*-linked ALS/FTD but not sporadic ALS. Curiously, the striatum harboured small punctate ubiquilin 2 aggregates in all cases examined, while large diffuse striatal ubiquilin 2 aggregates were specific to *UBQLN2*-linked ALS/FTD. Overall, ubiquilin 2 is mainly deposited in clinically unaffected regions throughout the CNS such that symptomology in *UBQLN2*-linked cases maps best to the aggregation of TDP-43.

## Introduction

A key pathological feature of amyotrophic lateral sclerosis (ALS) and frontotemporal dementia (FTD), present in 97% of ALS cases regardless of aetiology, is neuronal and oligodendrocytic aggregates of the TAR DNA binding protein of 43 kDa (TDP-43) [1–3]. In ALS caused by penetrant dominant genetic mutations, brain and/or spinal tissue often contains aggregates of both TDP-43 and the mutant-gene-encoded protein [4, 5]. Yet it is TDP-43 aggregate burden that correlates with neurodegeneration in sporadic ALS [6–8], and in *C9orf72*-linked ALS/FTD, which is characterised by additional deposition of dipeptide repeat proteins (DPRs) [9]. Whether the distribution of pathological TDP-43, or rather the mutant-gene-encoded protein, underpins clinical phenotype in cases with rarer genotypes is unknown. Mutations in *UBQLN2* cause X-linked dominant ALS with or without FTD [4]. *UBQLN2*-linked cases show deposition of TDP-43 and ubiquilin 2 aggregates in several CNS regions but these have not been mapped extensively, nor with respect to one another [4, 10–13].

Ubiquilin 2 is a ubiquitin-binding protein that delivers ubiquitinated substrates to the 26S proteasome for degradation [14, 15]. It also functions in autophagy, co-localising with LC3-II and optineurin in autophagosomes [16, 17], and regulating autophagosme acidification via the vacuolar ATPase proton pump [18]. Under cellular stress, and if not ubiquitin-bound, ubiquilin 2 undergoes liquid-liquid phase separation and is recruited to cytoplasmic stress granules [19]. The majority of ALS-causing mutations in ubiquilin 2 occur in or near a proline-rich repeat domain [4, 10, 12, 20–22], altering ubiquilin 2 phase separation dynamics, promoting liquid-to-solid transitions and, for certain mutants, driving ubiquilin 2 itself to aggregate [23]. Mutations in ubiquilin 2 also impair proteasome and/or autophagy function, thus promoting accumulation and aggregation of their degradation substrates including TDP-43 [4, 24, 25].

TDP-43 is a DNA- and RNA-binding protein with diverse roles in DNA maintenance and RNA processing; regulating DNA repair, mRNA stability and splicing, microRNA biogenesis and more [26, 27]. In most ALS cases, TDP-43 is found deposited in *post-mortem* brain and spinal cord tissue as neuronal and oligodendrocytic aggregates [1, 2, 28], predominantly in the motor cortex, spinal cord ventral horn, and cranial nerve nuclei [6]. Further ‘spreading’ of aggregates in ALS involves synaptically connected cortical and midbrain regions, with deposition of TDP-43 aggregates in the temporal lobe and hippocampus seen selectively in cases with abundant and distributed TDP-43 pathology (“stage 4”) [6].

Investigation of ubiquilin 2 and TDP-43 pathology in *UBQLN2*-linked ALS/FTD tissue has been limited, with ubiquilin 2-positive aggregates having only been identified in the hippocampus, spinal cord, neocortex, and brainstem [4, 10, 12, 13, 20, 29] (Fig. 1). In *UBQLN2*-linked ALS/FTD spinal cord, ubiquilin 2 aggregates have been shown to colocalise with TDP-43, however the same is also true of most non-*UBQLN2*-linked ALS/FTD [4, 12]. Our group recently showed that ubiquilin 2 in the hippocampus in *UBQLN2*-linked cases is more aggregation-prone than in non-*UBQLN2*-linked cases, with mutant but not wildtype ubiquilin 2 often aggregating independently of a known scaffold [13]. However, in other CNS regions in *UBQLN2*-linked disease, it is unclear whether ubiquilin 2 aggregation occurs together with TDP-43 or independently of a TDP-43 ‘scaffold’.

**Figure 1.**
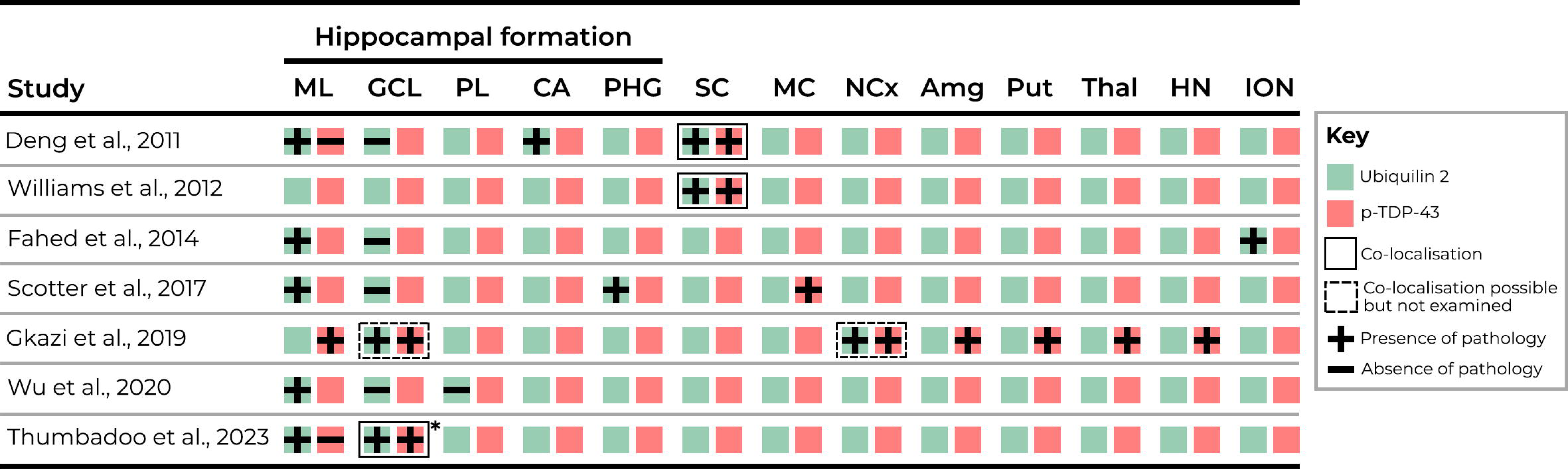
Ubiquilin 2 and TDP-43 labelling in previous studies largely fails to discriminate between independent aggregation and co-aggregation. Schematic summary of literature describing ubiquilin 2 and TDP-43 neuropathology in *UBQLN2*-linked human ALS/FTD. Ubiquilin 2 aggregates (green box with plus) or TDP-43 aggregates (red box with plus) in each CNS region were either; not both tested (one plus or minus), not both identified (one plus, one minus), both identified but co-localisation not tested (dashed black box), or co-localised (shared black box). *Co-localisation in certain cases only. Abbreviations: ML, molecular layer; GCL, granule cell layer; PL, polymorphic layer; CA, cornu ammonis; PHG, parahippocampal gyrus; SC, spinal cord; MC, motor cortex; NCx, neocortex; Amg, amygdala; Put, putamen; Thal, thalamus; HN, hypoglossal nucleus ION, inferior olivary nucleus.

A large multigenerational family with members in New Zealand, Australia, and the United Kingdom shows a c.1460C>T mutation in *UBQLN2*, encoding a threonine to isoleucine amino acid change (p.T487I), that co-segregates with ALS with or without FTD [12]. Donated brain and spinal cord tissue was available from three affected members of this family, each harbouring the p.T487I *UBQLN2* mutation. Here we employ multiplex immunohistochemistry to map ubiquilin 2 and TDP-43 aggregates across the CNS in these individuals, compared to controls and stage 4 sporadic ALS (sALS). We seek to characterise the differential aggregation propensities of mutant and wildtype ubiquilin 2 across the CNS, and to identify the CNS regions in which ubiquilin 2 aggregation occurs independently of TDP-43, to enable clinicopathological correlation and targeting of the true molecular ‘driver’ of pathogenesis.

## Materials and methods

### Systematic review of ubiquilin 2 and TDP-43 pathology in *UBQLN2*-linked ALS CNS tissue from published studies

Journal articles were identified using PubMed that performed immunohistochemical staining for ubiquilin 2 and/or TDP-43 in post-mortem ALS/FTD human brain tissue from cases with a *UBQLN2* mutation. Those that were not primary research articles or not published in English were excluded. Seven papers were identified from which neuropathological data for ubiquilin 2 and TDP-43 were extracted and tabulated (Figure 1).

### Patient demographics and CNS tissue

Formalin-fixed paraffin-embedded (FFPE) post-mortem brain tissue was obtained from eight neurologically normal control cases, three sporadic ALS (sALS) cases with phosphorylated TDP-43 proteinopathy (pTDP-43) classified as ‘stage 4’ according to Brettschneider’s neuropathological staging system [6], and three related *UBQLN2* p.T487I-linked familial ALS cases with or without FTD. Patient demographic and clinical information is summarised in Table S1. Tissue was obtained from up to 40 subregions of each brain, within the hippocampal formation and associated cortices, spinal cord, neocortex, striatum, medulla, substantia nigra, and cerebellum (Table S2 and S3). Tissue from all controls and sALS cases, and *UBQLN2*-linked ALS+FTD case MN17, was obtained from the Neurological Foundation Human Brain Bank at the Centre for Brain Research, Auckland, New Zealand, and processed as previously described [30]. It should be noted that the latter (MN17) was stored in fixative for 7 years before embedding. All NZ cases, but no controls, had confirmed pTDP-43 proteinopathy in the motor cortex. Tissue from two other *UBQLN2* p.T487I-linked cases were obtained from: the Victoria Brain Bank (pedigree ID V:7 in Figure 1A of [12], with ALS+FTD); and the Centre for Motor Neuron Disease Research, Macquarie University (pedigree ID IV:10 in Figure 1A of [12], with ALS). Clinical and neuropathological diagnoses were conducted as described previously [12, 29] or by the Victoria Brain Bank for case V:7 (diagnosis not previously described). Tissue was not always available for all regions for each case.

### Multiplexed fluorescent immunohistochemistry

Fluorescence immunohistochemistry was performed as described previously [31–33] using primary and secondary antibodies as described in Tables S4 and S5. The ubiquilin 2 antibody used here (mouse monoclonal anti-ubiquilin IgG_2a_, Santa Cruz #SC-100612) has been used by us previously for the detection of neuropathological aggregates [18, 29] and the manufacturer confirms is the same clone as that used by [4] and [12] (mouse monoclonal anti-ubiquilin IgG_2a_, clone 5F5, Novus Biologicals #H00029978-M03). *UBQLN2*-linked ALS+FTD case V:7 showed poor Hoechst labelling of nuclei, therefore a primary antibody against histone H3 was used as a nuclear marker. To aid detection of TDP-43 aggregates for quantification, we pooled pan- and phospho-TDP-43 antibodies (both rat IgG_2a_, identified only as ‘TDP-43’). Bleed-through (Fig. S1), secondaries-only, and neurologically normal control sections were included for each staining.

Cross-reactivity was observed during staining optimisation, between goat anti-mouse IgG_1_ secondary antibodies and rat IgG_2a_ primary antibodies. Therefore, for quantitative analysis of spinal cord and motor cortex sections which required labelling with both SMI-32 (mouse IgG_1_) and pooled pan- and phospho-TDP-43 antibodies (both rat IgG_2a_), two sequential rounds of immunofluorescent labelling were performed, as described previously [32].

### Image acquisition

Wide-field images for qualitative or quantitative analysis were acquired using a Zeiss Z2 Axioimager with a MetaSystems VSlide slide scanning microscope (20x dry magnification lens, 0.9 NA) with a Colibri 7 solid-state fluorescent light source. Filters were as described and validated previously [34]. Exposure times for each channel were set per case due to differences in immunoreactivity; *UBQLN2*-linked ALS+FTD case MN17 showed poor immunoreactivity overall, likely due to long-term fixation, therefore longer exposures were used. Confocal images were acquired using an Olympus FV1000 confocal microscope (100x magnification oil immersion lens, 1.4 NA, *Z*-step 0.4-1 µm) with FluoView 4.2 software (Olympus). Maximum intensity *Z*-projections were generated and processed using FIJI.

Final figures were compiled using FIJI software (v1.53c, National Institutes of Health) and Adobe Illustrator (Adobe Systems Incorporated, v24.3). Extracted single-channel images were imported into FIJI, merged, and pseudocoloured. For each channel, image intensity was adjusted in FIJI to best view the marker of interest, reduce background autofluorescence, and generate images representative of ubiquilin 2 and TDP-43 aggregate burden in each region.

### Qualitative analysis of ubiquilin 2 aggregates

To qualitatively assess which CNS regions showed ubiquilin 2 deposition independent of TDP-43, imaged sections were assessed for the abundance of ubiquilin 2-only (pTDP-43-negative) aggregates. Anatomical regions and sub-regions were delineated by anatomical markers as follows and as per Table S4: All sub-regions by neuronal marker MAP2; cortical layers and ventral horn motor neurons by SMI-32 [35, 36]; cerebellar cortical layers and dentate nucleus by calbindin [37, 38]; and globus pallidus externa, interna, and putamen by enkephalin [39, 40]. To account for the presence of fluorescent artefacts, ubiquilin 2-only aggregates were considered a consistent pathological feature of a subregion only when five or more aggregates were identified in that subregion.

### Quantitative analysis of ubiquilin 2-only, TDP-43-only, and ubiquilin 2/TDP-43 double immunopositive aggregates

Image analysis methods to quantify relative ubiquilin 2 and TDP-43 aggregate deposition were developed in MetaMorph v7.10.5.476 (Molecular Devices), similar to those previously described [8, 41]. Briefly, a composite image of MAP2 and Hoechst for each section was used to identify each subregion of interest and exclude tissue folds and defects. Nuclear TDP-43 (only detected in stage 4 sALS) was excluded by using the Count Nuclei application to segment all Hoechst-positive nuclei and removing the nuclear area from the TDP-43 image. Ubiquilin 2 and TDP-43 aggregates were then segmented using absolute thresholding by labelling intensity, excluding single positive pixels, and binary masks created. Thresholds were consistent within but not between cases. Binary masks were combined to create a ‘master mask’, within which each object was considered a single aggregate. Aggregates were then classified to be ubiquilin 2 or TDP-43 positive if the maximum intensity for that immunolabel was greater than the respective threshold used in the initial segmentation. This identified 3 aggregate populations: ubiquilin 2-only, TDP-43-only, and ubiquilin 2- and TDP-43-double-immunopositive. Aggregate densities were determined by normalising aggregate number to subregion area. Double-immunopositive aggregates were normalised to 1) total aggregates identified, 2) total ubiquilin 2-positive aggregates identified, and 3) total TDP-43-positive aggregates identified.

## Results

### Ubiquilin 2 and TDP-43 labelling in previous studies largely fails to discriminate between their independent aggregation and co-aggregation

Review of the literature describing ubiquilin 2 and TDP-43 neuropathology in *UBQLN2*-linked human ALS/FTD identified 7 results (Fig. 1). Ubiquilin 2 and TDP-43 neuropathology had been reported in many of the same CNS regions but only in hippocampus and spinal cord had co-immunolabelling been performed, including by us. Co-immunolabelling indicated that in hippocampus, mutant ubiquilin 2 aggregates either independently of TDP-43 (molecular layer [4, 13]) or co-aggregates with TDP-43 (granule cell layer [10, 13]), and in spinal cord, mutant ubiquilin 2 co-aggregates with TDP-43 [4, 12]. For other brain regions, the relationship between ubiquilin 2 and TDP-43 aggregation was unknown.

### Ubiquilin 2-only aggregates are seen across much of the CNS in *UBQLN2* p.T487I-linked ALS/FTD

To compare the distribution of aggregates of ubiquilin 2 with TDP-43 across the CNS, we compared tissue from 1-3 *UBQLN2* p.T487I-linked ALS/FTD cases with tissue from two stage 4 sALS cases and two non-neurodegenerative disease controls, from up to 40 CNS subregions (Table S2). We first identified aggregates positive for ubiquilin 2 and negative for TDP-43 (ubiquilin 2-only aggregates). Strikingly, ubiquilin 2-only aggregates comprised the vast majority of all ubiquilin 2 aggregates. They were clearly present (>5 aggregates per sub-region, exceeding the threshold of similar artefactual objects in control tissue) in every neocortical subregion; cervical, thoracic, and lumbar levels of the spinal cord; the dentate gyrus of the hippocampus (see also [13]); the subiculum, entorhinal cortex, and perirhinal cortex of the parahippocampus; most subregions of the medulla; the putamen, the ventral striatum, and the caudate nucleus. Ubiquilin 2-only aggregates were scarce in or absent from the *UBQLN2* cases in the pyramids, raphe nuclei, and medial lemniscus of the medulla; the globus pallidus; and the cerebellum. No ubiquilin 2 aggregates were identified in any region of the CNS of non-neurodegenerative disease control cases.

Confocal microscopy of aggregate morphology showed that ubiquilin 2 aggregates in the *UBQLN2* cases were predominantly small and punctate in almost all CNS subregions (Fig. 2). Unexpectedly given previous literature [4, 12], spinal cord lower motor neuron TDP-43 aggregates were ubiquilin 2-negative both in the *UBQLN2* cases and in stage 4 sALS (Fig. 2A, B). However, punctate ubiquilin 2-only aggregates were present in the spinal cord dorsal horn and motor cortex in the *UBQLN2* cases (Fig. 2C, E) while smaller dot-like ubiquilin 2-only aggregates were present in the spinal cord dorsal horn and motor cortex in stage 4 sALS cases (Fig. 2D, F).

**Figure 2.**
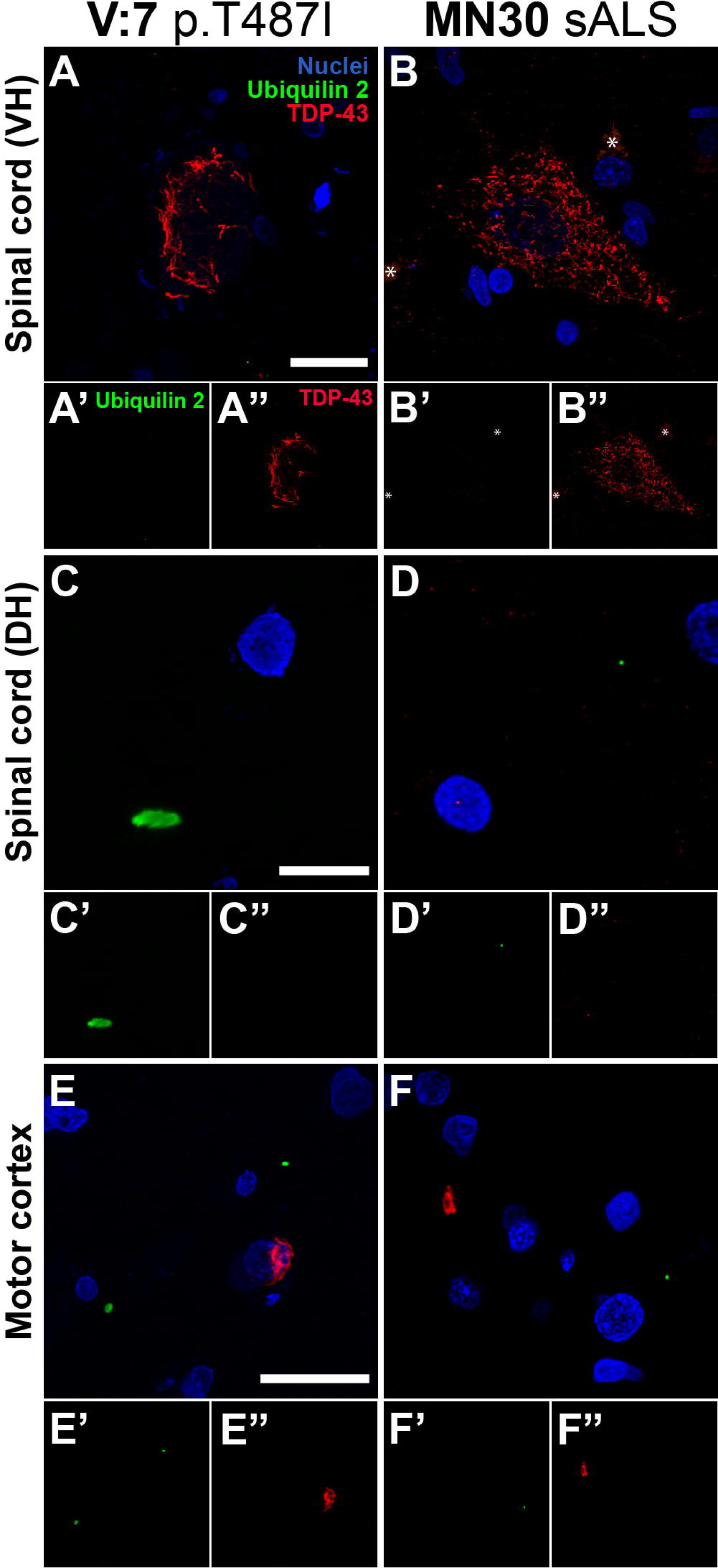
Morphology of ubiquilin 2 aggregates in *UBQLN2* p.T487I-linked ALS/FTD. Sections from across the CNS from representative *UBQLN2* p.T487I-linked ALS/FTD case V:7 (**A, C, E**) and representative stage 4 sALS case MN30 (**B, D, F**). Spinal cord ventral horn (VH) motor neurons demonstrated large TDP-43 aggregates (red) that were negative for ubiquilin 2 (green) in both *UBQLN2* p.T487I-linked ALS/FTD (**A**) and stage 4 sALS (**B**). Dorsal horn (DH) demonstrated small punctate ubiquilin 2 aggregates (green) that were negative for TDP-43 (red) in *UBQLN2* p.T487I-linked ALS/FTD (**C**) and in stage 4 sALS where they were smaller and dot-like (**D**). Motor cortex demonstrated small punctate ubiquilin 2 aggregates (green) that were negative for TDP-43 (red) in *UBQLN2* p.T487I-linked ALS/FTD (**E’**) and in stage 4 sALS where they were dot-like (**F’**); and TDP-43 aggregates (red) that were negative for ubiquilin 2 (green) in both *UBQLN2* p.T487I-linked ALS/FTD (**E”**) and stage 4 sALS (**F”**). Scale bars 20 µm.

**Figure 3.**
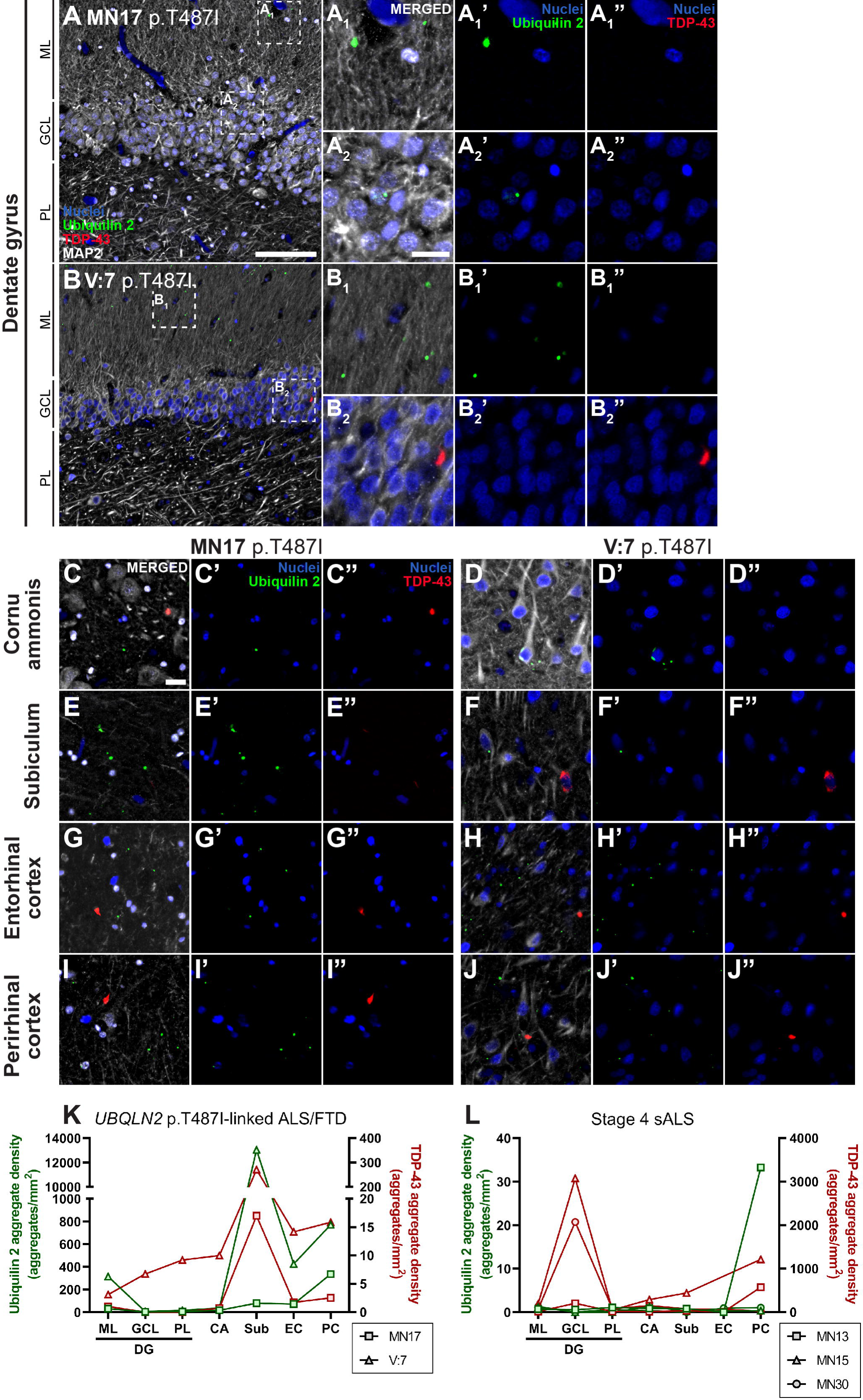
Ubiquilin 2-only and TDP-43-only aggregates in the *UBQLN2* p.T487I-linked ALS/FTD hippocampal formation, and entorhinal and perirhinal cortices. Hippocampal sections from 2 *UBQLN2* p.T487I-linked ALS/FTD cases (**A-J”**) demonstrated punctate aggregates in the dentate gyrus molecular layer (**A_1_, B_1_**) that were positive for ubiquilin 2 only (**A_1_’, B’_1_,** green) and negative for TDP-43 (**A_1_”, B_1_”**). Case V:7 showed rare granule cell layer cytoplasmic aggregates of TDP-43 only (**B_2_”,** red) that were negative for ubiquilin 2 (**B_2_’**). Cornu ammonis and parahippocampal subregions showed abundant punctate ubiquilin 2-only aggregates (**C’-J’**) and fewer TDP-43-only aggregates (**C”-J”**) that were each negative for the other protein. Quantification of the density of ubiquilin 2-only (green) and TDP-43-only (red) aggregates showed that in *UBQLN2* p.T487I-linked ALS/FTD cases both aggregate types were highest in subiculum (**K**), while in stage 4 sALS ubiquilin 2-only aggregates were rare or absent but TDP-43-only aggregates were found throughout the hippocampal formation, particularly in the granule cell layer (**L**). Scale bars in A, B, 100 µm; others 20 µm. Abbreviations: ML, molecular layer; GCL, granule cell layer; PL, polymorphic layer; CA, cornu ammonis; Sub, subiculum; EC, entorhinal cortex; PC, piriform cortex.

Because we did not observe ubiquilin 2 co-labelling of TDP-43 aggregates in the spinal cord ventral horn, we subsequently validated our monoclonal anti-ubiquilin 2 antibody (#SC-100612, clone 5F5, IgG_2a_). Labelling using the same 5F5 ubiquilin 2 antibody clone, but purchased from Novus Biologicals, at the recommended concentration (#H00029978-M03, clone 5F5, IgG_2a_ 1:250) showed occasional detection of dense TDP-43 aggregates in the granule cell layer, but also high background and non-specific (nuclear) labelling (data not shown). We therefore co-labelled the Santa Cruz 5F5 clone with a different Novus Biologicals monoclonal anti-ubiquilin 2 antibody (#NBP2-25164, clone 6H9, IgG_1_) which both showed the same pattern of labelling in the *UBQLN2* and stage 4 sALS spinal cord (Fig. S2, S3): TDP-43 aggregates in the ventral horn were ubiquilin 2-negative irrespective of the ubiquilin 2 antibody used (Fig. S2E, F, S3A, B), and ubiquilin 2-only inclusions in the dorsal horn were identified with both ubiquilin 2 antibodies (Fig. S2A). This was consistent with observations in the hippocampus: TDP-43 aggregates in the granule cell layer were ubiquilin 2-negative irrespective of the ubiquilin 2 antibody used (Fig. S2K, L, S3C, D), and ubiquilin 2-only aggregates in the molecular layer were identified by both ubiquilin 2 antibodies (Fig. S2G, H, I). We were therefore confident that the monoclonal anti-ubiquilin 2 antibody from Santa Cruz was accurately detecting ubiquilin 2 aggregates in the CNS and was appropriate for use in subsequent experiments.

### Ubiquilin 2-only aggregates, but not TDP-43-only aggregates, are abundant in *UBQLN2* p.T487I-linked ALS/FTD hippocampal formation, and entorhinal and perirhinal cortices

Having qualitatively identified CNS regions in *UBQLN2* p.T487I-linked ALS/FTD with deposition of ubiquilin 2-only aggregates, we next conducted automated quantitative analysis in 2-3 *UBQLN2* p.T487I-linked ALS/FTD cases compared to 3 stage 4 sALS cases, in selected regions containing ubiquilin 2 aggregates (Table S3). As noted, the middle and lower medulla and substantia nigra contained ubiquilin 2-only aggregates, but these were not quantified (Fig. S4, S5, S6).

**Figure 4.**
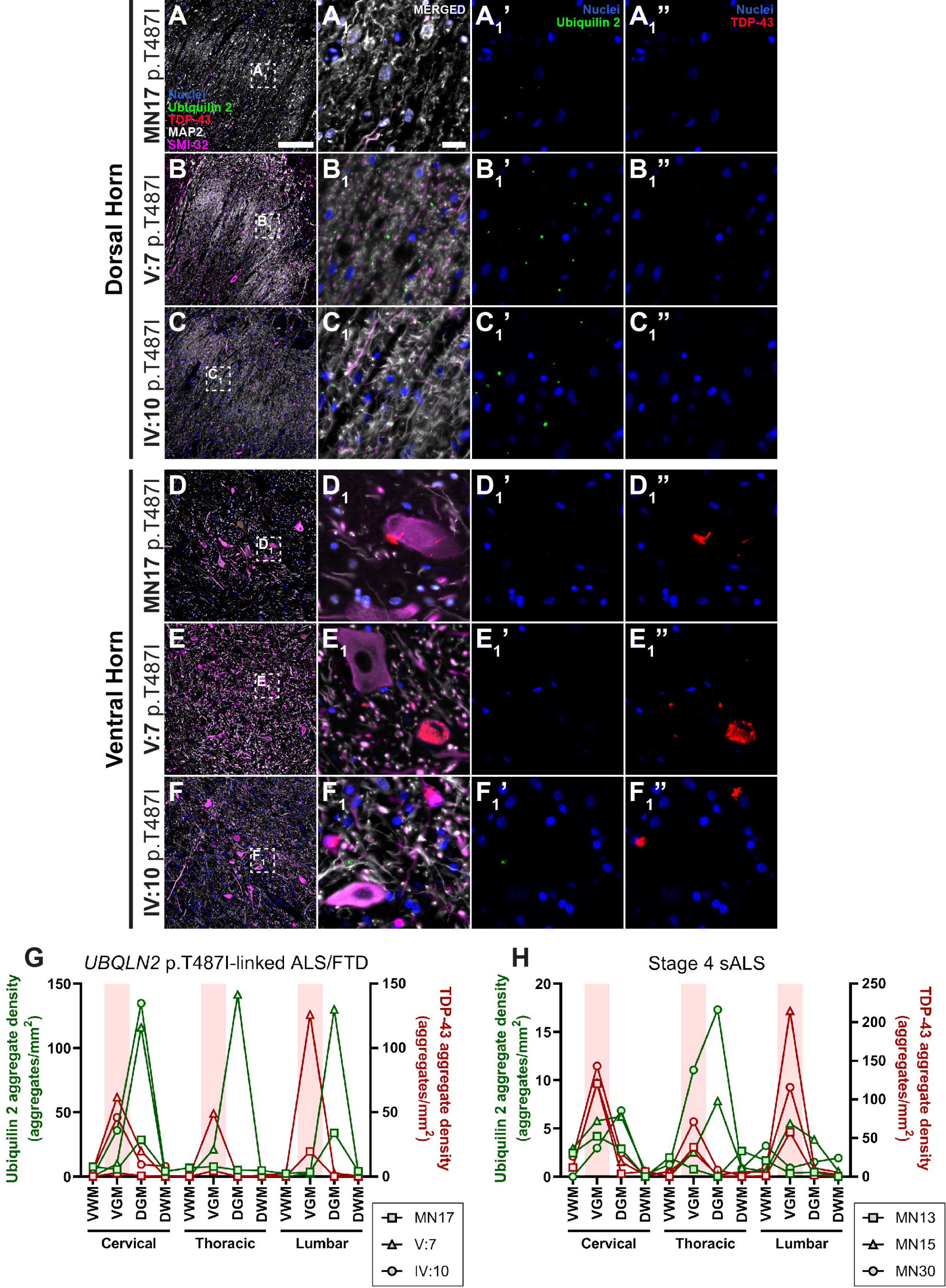
Ubiquilin 2-only and TDP-43-only aggregates in the *UBQLN2* p.T487I-linked ALS/FTD spinal cord. Spinal cord sections from 3 *UBQLN2* p.T487I-linked ALS/FTD cases (**A-F_1_”**) demonstrated punctate aggregates in the dorsal horn (**A, B, C**) that were positive for ubiquilin 2 only (**A_1_’, B_1_’, C_1_’,** green) and negative for TDP-43 (**A_1_”, B_1_”, C_1_”**). The ventral horns showed abundant cytoplasmic and neuropil TDP-43-only aggregates (**D_1_”, E_1_”, F_1_”,** red) that were negative for ubiquilin 2 (**D_1_’, E_1_’, F_1_’**). Quantification of the density of aggregates confirmed that in *UBQLN2* p.T487I-linked ALS/FTD cases ubiquilin 2-only aggregates (green) were most abundant in the dorsal horn but TDP-43-only aggregates (red) were most abundant in the ventral horn (highlighted by pale red shading) (**G**). The same pattern was observed in stage 4 sALS cases but with only rare ubiquilin 2-only aggregates in the dorsal horn (**H**). Scale bar in main images, 200 µm; in insets 20 µm. Abbreviations: VWM, ventral white matter; VGM, ventral grey matter; DGM, dorsal grey matter; DWM, dorsal white matter.

**Figure 5.**
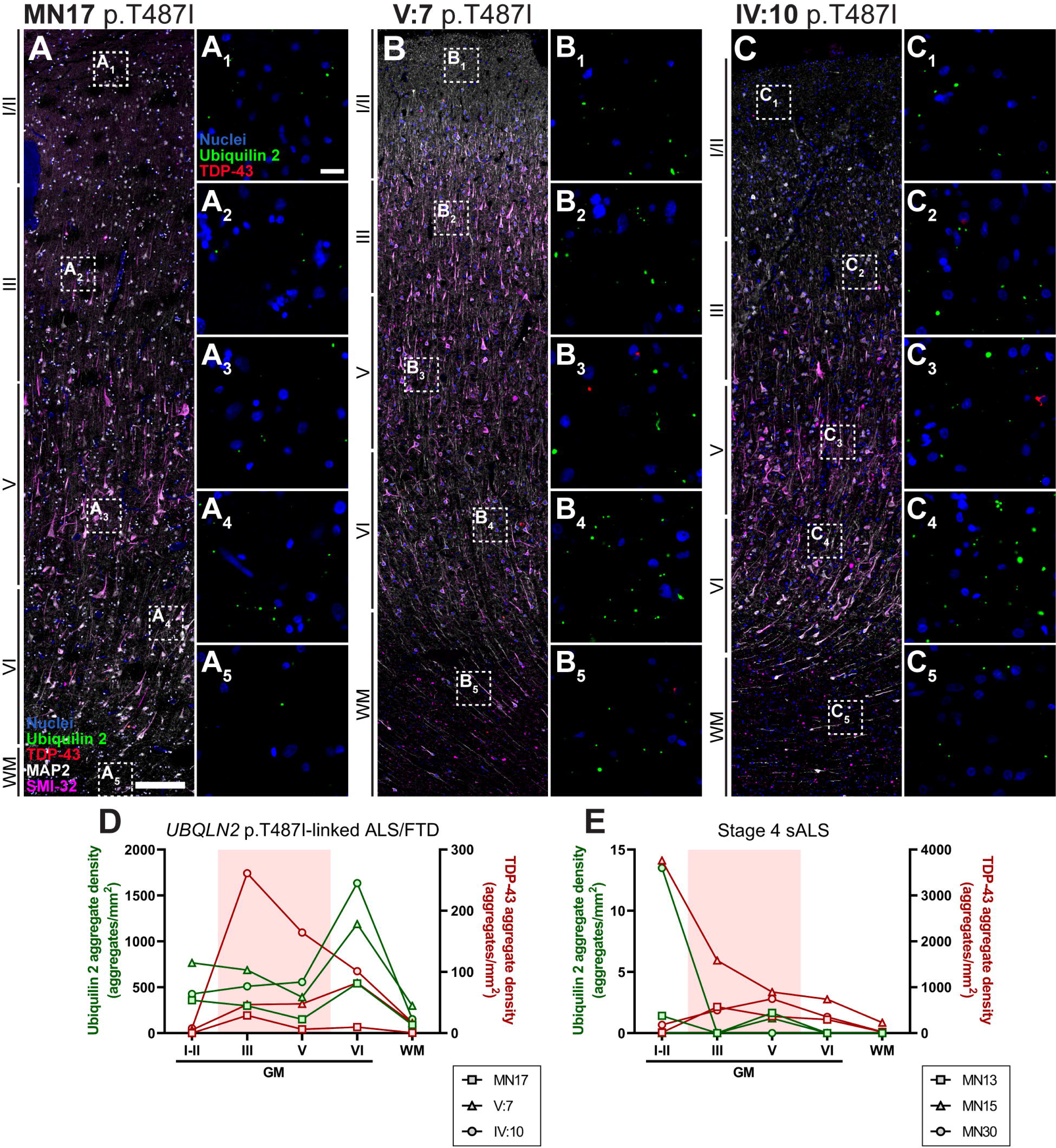
Ubiquilin 2-only and TDP-43-only aggregates in the *UBQLN2* p.T487I-linked ALS/FTD motor cortex. Motor cortex sections from 3 *UBQLN2* p.T487I-linked ALS/FTD cases (**A-C_5_**) demonstrated aggregates in the cortical layers I-VI that were positive for ubiquilin 2 only (green) or TDP-43 only (red), but rarely positive for both (**A_1_-A_5_, B_1_-B_5_, C_1_-C_5_**). Quantification of the density of aggregates demonstrated that in *UBQLN2* p.T487I-linked ALS/FTD cases ubiquilin 2-only aggregates (green) were most abundant in layer VI, but TDP-43-only aggregates (red) were found throughout the cortical layers with case IV:10 showing higher density in layers III and V (highlighted by pale red shading) (**D**). Stage 4 sALS cases showed very rare ubiquilin 2-only aggregates but TDP-43-only aggregates were found throughout the cortical layers (**E**). Scale bar in main images, 200 µm; in insets 20 µm. Abbreviations: GM, grey matter; WM, white matter.

**Figure 6.**
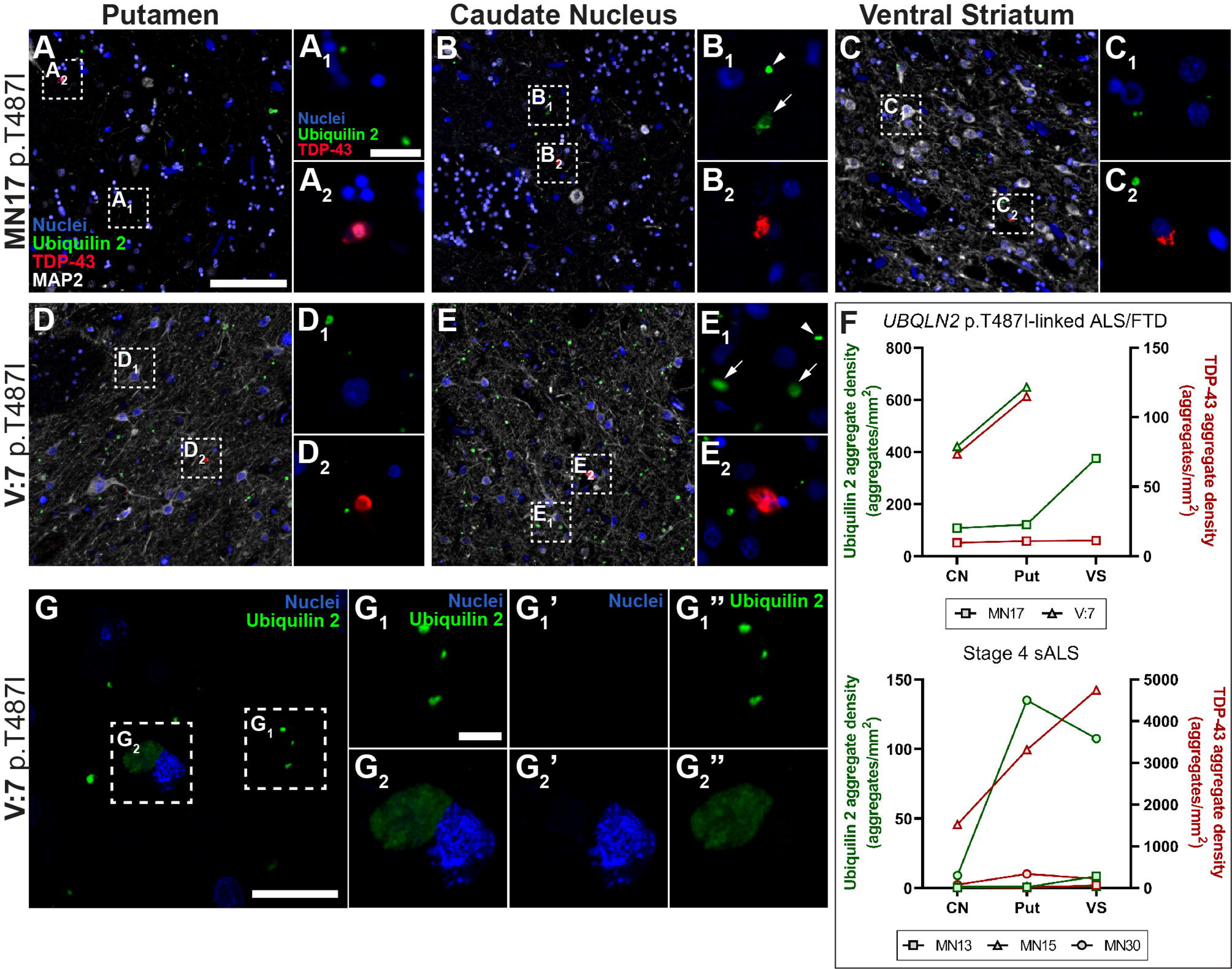
Ubiquilin 2-only and TDP-43-only aggregates in the *UBQLN2* p.T487I-linked ALS/FTD striatum. Striatal sections from 2 *UBQLN2* p.T487I-linked ALS/FTD cases demonstrated abundant punctate and some large diffuse aggregates in the putamen, caudate nucleus, and ventral striatum, that were positive for ubiquilin 2 only and negative for TDP-43 (**A-E_2_**, green). TDP-43-only aggregates were also present in these subregions but were less abundant (**A-E_2_**, red). Quantification of the density of aggregates confirmed that in *UBQLN2* p.T487I-linked ALS/FTD cases ubiquilin 2-only aggregates (green) were more abundant than TDP-43-only aggregates (red) (**F, upper**). Stage 4 sALS case MN30 showed deposition of ubiquilin 2-only aggregates in putamen and ventral striatum, while case MN15 showed particularly abundant TDP-43-only aggregates across the striatum (**F, lower**). Confocal imaging of the caudate nucleus (**G**) showed many small punctate ubiquilin 2 aggregates (green, **G_1_**) and few large diffuse ubiquilin 2 aggregates (green, **G_2_**). Scale bar in main images A-E, 100 µm; in insets of A-E and in image G, 20 µm; in insets of G, 5 µm. Abbreviations: CN, caudate nucleus; Put, putamen; VS, ventral striatum.

In the hippocampal formation, ubiquilin 2-only aggregates in the *UBQLN2* cases were found in the molecular layer, subiculum, entorhinal cortex, and perirhinal cortex (Fig. 3). As we have shown previously [13], ubiquilin 2-only aggregates in *UBQLN2* p.T487I-linked ALS/FTD were rare in or absent from the granule cell layer or cornu ammonis (Fig. 3). Ubiquilin 2-only aggregates in stage 4 sALS were generally absent from the hippocampal formation and associated cortices, with their highest density being in the perirhinal cortex of case MN13 where they were primarily dot-like (Fig. S8, 3L). Occasionally, larger ubiquilin 2 aggregates that colocalised with TDP-43 were found in stage 4 ALS (Fig. S7).

In contrast, TDP-43-only aggregates in the *UBQLN2* cases were found throughout the hippocampus at low density, and in the subiculum at moderate density (Fig. 3). TDP-43-only aggregates in stage 4 sALS were also found throughout the hippocampus and subiculum and were particularly abundant in the granule cell layer (Fig. 3L, S8), as reported previously [6]. Overall, hippocampal ubiquilin 2-only aggregates were predominantly seen in *UBQLN2* p.T487I cases, and TDP-43-only aggregates in stage 4 sALS cases, with a regional relationship between ubiquilin 2 and TDP-43 emerging in *UBQLN2* cases in the subiculum.

### Ubiquilin 2-only aggregates are abundant in *UBQLN2* p.T487I-linked ALS/FTD dorsal horn, but not ventral horn, of the spinal cord

In the spinal cord, ubiquilin 2-only aggregates were found almost exclusively in the dorsal grey matter (particularly layer II) in the *UBQLN2* cases in which they were abundant (Fig. 4), and stage 4 sALS in which they were scarce (Fig. S9, 4H). This was true for cervical, thoracic, and lumbar segments of the cord.

In contrast, TDP-43-only aggregates were found predominantly in the ventral grey matter in the *UBQLN2* cases (Fig. 4), and stage 4 sALS (Fig. S9, 4H). This was true for cervical, thoracic and lumbar segments. TDP-43-only aggregates were rare in or absent from the white matter and dorsal grey matter in the *UBQLN2* cases (Fig. 4) and stage 4 sALS (Fig. S9, 4H). As described above, and validated with a different monoclonal anti-ubiquilin 2 antibody (Fig. S2, S3), TDP-43 aggregates in the spinal cord did not co-label with ubiquilin 2 in any case. Overall, spinal cord ubiquilin 2-only aggregates were predominantly found in *UBQLN2* cases dorsally, but TDP-43-only aggregates were seen in both *UBQLN2* cases and stage 4 sALS cases ventrally. Again, there was a lack of regional relationship between aggregates of the two proteins.

### Ubiquilin 2-only aggregates in *UBQLN2* p.T487I-linked ALS/FTD are abundant throughout the motor cortex

In the motor cortex, ubiquilin 2-only aggregates in *UBQLN2* p.T487I-linked ALS/FTD were found throughout the cortical layers, being particularly abundant in layer VI (Fig. 5). Ubiquilin 2-only aggregates in stage 4 sALS were rare or absent from the entire motor cortex, with their highest density being in layers I/II of case MN30 (Fig. S10, 5E).

TDP-43-only aggregates in the *UBQLN2* cases were also found throughout the cortical layers, but at lower density than ubiquilin 2, and mostly driven by ALS case IV:10 (Fig. 5). TDP-43-only aggregates in stage 4 sALS were found throughout the cortical layers with their highest density being in layers I/II of case MN15 (Fig. S10, 5E). As seen in hippocampus *and* spinal cord, motor cortex ubiquilin 2-only aggregates were predominantly seen in the *UBQLN2* cases, while, as seen in hippocampus, TDP-43-only aggregates were predominantly seen in stage 4 sALS cases, with a lack of regional relationship.

### Ubiquilin 2-only aggregates in *UBQLN2 p.*T487I-linked ALS/FTD are abundant throughout the striatum and can also be found in stage 4 sALS striatum

In the striatum, ubiquilin 2-only aggregates in *UBQLN2* p.T487I-linked ALS/FTD were detected throughout the caudate nucleus, putamen, and ventral striatum, however tissue was only available for 1-2 cases (Fig. 6). Ubiquilin 2-only aggregates in stage 4 sALS were rare but present in cases MN13 and MN15 and more abundant in case MN30 (Fig. S11, 6G). When ubiquilin 2 aggregates were found in stage 4 sALS striatum, they occasionally co-labelled with TDP-43 (Fig. S7D). In addition to the small punctate ubiquilin 2-only aggregates also described in other regions of the CNS, larger diffuse ubiquilin 2-only aggregates were unique to the striatum (Fig. 6B_1_, E_1_ arrows, G_2_).

TDP-43-only aggregates were moderately abundant in striatum in *UBQLN2* case V:7 but less so in MN17 (Fig. 6). TDP-43-only aggregates in stage 4 cases were abundant in ALS/FTD case MN15 and present at low to medium density in sALS cases MN13 and MN30 (Fig. S11, 6F). Overall, the striatum showed highly variable pathology but emerged as a region with unique large and diffuse inclusions of ubiquilin 2 in *UBQLN2* p.T487I-linked cases. Striatum was also the only brain region with substantial deposition of ubiquilin 2-only aggregates in a stage 4 sALS case, albeit mainly one case.

### Ubiquilin 2- and TDP-43-double immunopositive aggregates are differentially distributed in the CNS between *UBQLN2 p.*T487I-linked ALS/FTD and stage 4 sALS

As described in results above, we examined aggregates ‘double immunopositive’ for ubiquilin 2 and TDP-43 (Fig. S7). Double-immunopositive aggregates in a given case were always rare as a proportion of total aggregates, achieving their highest proportions in the *UBQLN2* cases in the striatum (up to ∼4% of total aggregates, mainly of the large diffuse kind unique to striatum) (Fig. S7C). Double-immunopositive aggregates accounted for up to ∼70% in the hippocampus, and up to ∼25% in the striatum, of the total rare ubiquilin 2 aggregates of sALS cases, suggesting that TDP-43 can occasionally drive wildtype ubiquilin 2 aggregation in sALS (Fig. S7D). Conversely, very few of the abundant total ubiquilin 2 aggregates in the *UBQLN2* cases were TDP-43 positive, as already noted (Fig. S7D). Conversely, approximately one third of the total rare TDP-43 aggregates were ubiquilin 2 positive (Fig. S7E), together suggesting that mutant ubiquilin 2 is intrinsically aggregation-prone and does not require, yet may seed, TDP-43 aggregation.

## Discussion

Most ALS and many FTD cases are characterised by TDP-43 aggregation [2, 42], which causes neurotoxicity through loss or gain of TDP-43 function [43–45]. Much research effort has been dedicated to determining whether aggregation of other proteins in ALS and FTD is similarly neurotoxic, or whether TDP-43 is the driver of neurodegeneration. The patterning of TDP-43 aggregate deposition in sporadic ALS, being most severe in the upper and lower motor neurons of the motor cortex and spinal cord, respectively, reflects the major symptomology as a paralysing motor disorder [6, 7]. Determining the patterning of other aggregating proteins with respect to neurodegeneration, symptomology, and the pattern of TDP-43 aggregation, could therefore reveal their relative contribution to disease. We mapped ubiquilin 2 and TDP-43 aggregates across the CNS in *UBQLN2* p.T487I-linked ALS with or without FTD, finding that mutant ubiquilin 2 aggregates were abundant throughout much of the CNS; predominantly in different CNS subregions from and not colocalised with TDP-43. The broad distribution of ubiquilin 2 aggregates in clinically unaffected regions in these *UBQLN2*-linked cases supports the hypothesis that the aggregation of TDP-43 drives neurodegeneration more quickly or potently than that of ubiquilin 2.

### Regional distribution of ubiquilin 2- and/or TDP-43-containing aggregates in *UBQLN2*-linked ALS/FTD

Regions of ubiquilin 2 aggregation described in previous studies of *UBQLN2*-linked ALS/FTD were spread across the CNS, including the neocortex, hippocampal formation, brain stem, and spinal cord [4, 10, 12, 13, 18, 20, 29]. Here we examined 40 brain subregions across the CNS, finding ubiquilin 2-only aggregates across the neocortex and enriched in layer VI of the motor cortex, in the dorsal and rarely the ventral horns of the spinal cord, in the molecular layer of the dentate gyrus of the hippocampus, the substantia nigra, and throughout the hippocampal-associated cortices, medulla, and striatum. It should be noted that tissue from only cases with the *UBQLN2* p.T487I mutation were analysed, to limit heterogeneity and discern clear patterns of aggregate distribution associated with this variant. An important future direction will be to determine whether these aggregate distribution patterns extend to cases with other *UBQLN2* mutations.

The vast majority of ubiquilin 2-only aggregates were punctate and neuritic or found in the neuropil, with larger diffuse aggregates detected mainly in the ventral striatum. Ubiquilin 2-only aggregates occasionally localised to MAP2-positive neurites and cell bodies. The majority of aggregates independent of MAP2-positive structures may have localised to astrocytes, microglia, or to MAP2-negative neuronal subtypes [46] or axons, given that MAP2 strongly labels dendrites but not axons [47, 48].

Of the regions we identified with aggregates of ubiquilin 2 in *UBQLN2*-linked ALS/FTD, the motor cortex, striatum, substantia nigra, and most subregions of the medulla had not previously been reported, while the hippocampal molecular layer had been reported in almost all previous studies. Notably, while the hippocampal molecular layer *did* show dense ubiquilin 2 aggregates in these cases, their density was 2-40x higher in the subiculum and perirhinal cortex. Compared to the hippocampal molecular layer, ubiquilin 2 aggregate density was also higher in the striatum and layer VI of the motor cortex. The focus upon the hippocampus as the major CNS region in which mutant ubiquilin 2 aggregates should therefore be reconsidered.

### Is hippocampal ubiquilin 2 aggregation a determinant of frontotemporal dysfunction?

Overall, hippocampal ubiquilin 2-only aggregates were predominantly seen in *UBQLN2* p.T487I cases, and hippocampal TDP-43-only aggregates in stage 4 sALS cases. The focus on hippocampal ubiquilin 2 aggregates in previous literature regarding *UBQLN2*- linked ALS/FTD may derive from their appearance also in *C9orf72*-linked ALS/FTD, but not in other forms of ALS [4, 49]. It is tempting to speculate that hippocampal ubiquilin 2 aggregates underpin the high frequency of co-morbid FTD in both *UBQLN2*-linked and *C9orf72*-linked cases [42, 50, 51]. FTD is characterised by neuronal loss beginning in the frontal and temporal lobes including the hippocampus, but progressing to involve most of the cortex and parts of the subcortex [52, 53]. While we found moderate-to-high deposition of ubiquilin 2 in the hippocampus in two *UBQLN2*-linked ALS+FTD cases, no hippocampal tissue was available from the *UBQLN2*-linked pure ALS case V:10, precluding even rudimentary analysis of a link between hippocampal ubiquilin 2 aggregation and FTD phenotype. Certainly, hippocampal deposition of TDP-43 (in the granule cells) is a known determinant of frontotemporal dysfunction, seen in ∼30% of stage 4 or *C9orf72*-linked ALS [6] or *UBQLN2*-linked ALS/FTD [13].

### Is motor region ubiquilin 2 aggregation a determinant of motor dysfunction?

In both the spinal cord and motor cortex, ubiquilin 2-only aggregates were predominantly seen in *UBQLN2* p.T487I cases, while TDP-43-only aggregates were predominantly seen in stage 4 sALS cases, with a lack of regional relationship. TDP-43 aggregates were found in the somata of motor neurons known to degenerate in ALS/FTD [6, 7]; the Betz cells of layer Vb of the motor cortex and the lower motor neurons of layer IX of the ventral spinal cord. In contrast, ubiquilin 2-only aggregates were deposited in layer VI of the motor cortex and layer II of the dorsal horn of the spinal cord. So does ubiquilin 2 aggregation promote or act as a biomarker of motor dysfunction, or is it simply an epiphenomenon? Motor cortex layer VI deposition of ubiquilin 2 is intriguing, given that the Betz cells are instead localised to the lower part of layer V, termed layer Vb [54]. Layer VI is the most complex of the cortical layers, containing a wide variety of neuronal cell types including excitatory pyramidal neurons that project dendrites and axons to layer V, establishing reciprocal connectivity [55–57]. However, ubiquilin 2-only aggregates in layer VI are punctate and co-localise frequently with MAP2-positive and occasionally with SMI-32-positive neurites, therefore are likely to be neuritic or localised to the neuropil rather than somatic. It may be that layer VI ubiquilin 2-only aggregates lie in neurites that derive from neurons elsewhere; Betz cells themselves project dendrites to layer VI and may be the source of layer VI ubiquilin 2-only aggregates [54].

In the spinal cord, our findings were wholly unexpected. First, we detected abundant punctate ubiquilin 2-only aggregates in the dorsal horn in *UBQLN2*-linked ALS/FTD cases that were scarce in sALS. The dorsal horn relays sensory inputs, and is generally affected only subtly by ALS, if at all [58–61]. The pathomechanisms and consequences of deposition of these aggregates warrant further investigation. Second, ubiquilin 2 has previously been shown to co-label TDP-43 aggregates in ventral horn lower motor neuron somata in *UBQLN2*-linked ALS/FTD cases and in sALS [4, 12], yet we never saw this even with striking TDP-43 aggregate burden in the ventral horn and abundant ubiquilin 2-labelling in the dorsal horn. As described in Materials and Methods, the antibody we used is the same clone as that used by those studies. We have tested this antibody at a high concentration (2 µg/mL), as well as another ubiquilin 2 monoclonal antibody (Novus Biologicals NBP2-25164), but we do not detect ubiquilin 2 co-labelling of TDP-43 aggregates in ventral horn in either *UBQLN2*-linked ALS/FTD cases or in sALS. If indeed ubiquilin 2 does not accumulate in ventral horn motor neurons, the main driver of lower motor neuron loss in *UBQLN2*-linked ALS may be deposition and dysfunction not of ubiquilin 2 but of TDP-43.

### Unique ubiquilin 2 pathology in the striatum

The striatum was unusual for the abundance of larger diffuse ubiquilin 2 aggregates that had the globular appearance of neuronal cytoplasmic inclusions. We found previously that ubiquilin 2 decorates cytoplasmic inclusions of TDP-43 induced by autophagy and/or proteasome blockade *in vitro* [62], yet in *UBQLN2* p.T487I-linked ALS/FTD these inclusions were devoid of TDP-43. We hypothesise that only in the striatum does mutant ubiquilin 2 aggregate significantly in the cell soma, in contrast to its neuritic or neuropil aggregation elsewhere in the CNS.

### Co-aggregation versus independent aggregation of ubiquilin 2

This brings us to our final discussion point, regarding the enhanced aggregation propensity of mutant compared to wildtype ubiquilin 2. This is indicated by i) the abundance of ubiquilin 2 aggregates in *UBQLN2* p.T487I cases compared even to stage 4 sALS cases with high TDP-43 aggregate loads; ii) the presence of large ubiquilin 2 neuronal cytoplasmic inclusions only in p.T487I cases; and most notably, iii) the fact that nearly all ubiquilin 2 aggregates in *UBQLN2* p.T487I cases are negative for TDP-43 whereas rare ubiquilin 2 aggregates in stage 4 sALS often incorporate TDP-43. We propose that TDP-43 drives the infrequent aggregation of wildtype ubiquilin 2 in sALS, while mutant ubiquilin 2 is intrinsically aggregation-prone and requires no driver.

## Conclusion

Here we demonstrate that the ubiquilin 2 aggregates described in previous papers in *UBQLN2*-linked ALS/FTD are found across most of the CNS, where they are punctate in the neurites and neuropil. A unique population of larger diffuse ubiquilin 2 aggregates are also found in the striatum, but both punctate and diffuse ubiquilin 2 aggregates are generally negative for TDP-43. Overall symptomology in *UBQLN2*-linked ALS/FTD maps best to the aggregation of TDP-43.

## List of abbreviations

ALS: Amyotrophic lateral sclerosis
*C9orf72*: Chromosome 9 open reading frame 72
CNS: Central nervous system
DPR: Dipeptide repeat
FFPE: Formalin-fixed paraffin-embedded
FTD: Frontotemporal dementia
PBS: Phosphate-buffered saline
pTDP-43: Phosphorylated TDP-43
sALS: Sporadic ALS
TDP-43: Transactive response DNA binding protein 43 kDa
*UBQLN2*: Ubiquilin 2 gene

## Declarations

### Ethics approval and consent to participate

All protocols were approved by the University of Auckland Human Participants Ethics Committee (New Zealand) and carried out as per approved guidelines. This study was also approved by the Human Research Ethics Committee of Macquarie University (520211013428875) and the Concord Hospital Research Ethics Committee.

### Consent for publication

Informed donor consent was obtained at each site.

### Availability of data and material

The datasets used and/or analysed during the current study are available from the corresponding author on reasonable request.

### Competing interests

The authors declare that they have no competing interests.

## Funding

KT was supported by a doctoral scholarship from Amelia Pais-Rodriguez and Marcus Gerbich. HCM was supported by a Health Education Trust Postdoctoral Fellowship. ELS was supported by Marsden FastStart and Rutherford Discovery Fellowship funding from the Royal Society of New Zealand [15-UOA-157, 15-UOA-003]. This work was also supported by grants from Motor Neuron Disease NZ, Freemasons Foundation of New Zealand, Matteo de Nora, and PaR NZ Golfing. No funding body played any role in the design of the study, nor in the collection, analysis, or interpretation of data nor in writing the manuscript.

## Authors’ contributions

LRN, KMT, BVD, HCM, MEVS conducted or designed experiments; LRN, MEVS performed data analysis and designed analysis methods; CT and CM conducted neuropathological diagnostics; SY, IPB, RLMF, MAC, FH, GAN coordinated the banking and use of human tissue for study; LRN, KMT, MEVS, ELS wrote the manuscript; ELS conceived of and designed the study. All authors read, edited, and approved the final manuscript.

## Acknowledgements

This publication is dedicated to the incredible patients and families who contribute to our research. We thank Marika Eszes at the Centre for Brain Research, University of Auckland, New Zealand, and the Neurological Foundation of New Zealand for their ongoing financial support of the Human Brain Bank. We also thank Fairlie Hinton at the Victorian Brain Bank, which is supported by The Florey, The Alfred, Victorian Institute of Forensic Medicine and Coroners Court of Victoria and funded in part by Fight Parkinson’s, MND Victoria, FightMND, Yulgilbar Foundation and Ian and Maria Cootes. The imaging data reported in this paper were obtained at the Biomedical Imaging Research Unit, operated by the Faculty of Medical and Health Sciences’ Technical Services at the University of Auckland.

## Notes

### Competing Interest Statement

The authors have declared no competing interest.

### Summary of Updates

Minor updates to all figures and manuscript.

