## Supplementary material for "Distribution of ubiquilin 2 and TDP-43 aggregates throughout the CNS in *UBQLN2* p.T487I-linked amyotrophic lateral sclerosis and frontotemporal dementia": Fig. S

### Supplementary Tables

Table S1. Control and ALS patient demographic and clinical information.

| Case type | UoA case code | Age at onset (y) | Age at death (y) | Sex | PMD (h) | Clinical presentation or cause of death | Genetic mutation |
| --- | --- | --- | --- | --- | --- | --- | --- |
| Neurologically normal control | H211 | N/A | 41 | M | 8 | Ischaemic heart disease | N/A |
|  | H215 | N/A | 67 | F | 23.5 | Ischaemic heart disease | N/A |
|  | H231 | N/A | 65 | M | 8 | Ischaemic heart disease | N/A |
|  | H238 | N/A | 63 | F | 16 | Dissecting aortic aneurysm | N/A |
|  | H239 | N/A | 64 | M | 15.5 | Ischemic Heart Disease | N/A |
|  | H243 | N/A | 77 | F | 13 | Ischaemic heart disease- coronary atherosclerosis | N/A |
|  | H245 | N/A | 63 | M | 20 | Asphyxia | N/A |
|  | H248 | N/A | 98 | M | 10.5 | Congestive heart failure | N/A |
| sALS | MN12 | 46 | 49 | M | 34 | ALS | None found^a^ |
|  | MN13 | 55 | 55 | M | 10 | ALS | None found^a^ |
|  | MN15 | 52 | 54 | F | 18 | ALS + FTD | None found^a^ |
|  | MN30 | 84 | 84 | F | 17.5 | ALS | N/D |
| *UBQLN2*-linked ALS | MN17 | 58 | 61 | F | 69 | ALS + FTD | *UBQLN2* p.T487I |
|  | IV:10 | 34 | 35 | M | ^b^ | ALS | *UBQLN2* p.T487I |
|  | V:7 | 29^c^ | 39 | M | 25.5 | ALS + FTD | *UBQLN2* p.T487I |

N/A, not applicable. N/D, not done.

^a^ Negative for mutations in *C9orf72*, *TARDBP*, *SOD1* and *FUS*, as tested in Scotter et al. (2017).

^b^ Information not available.

^c^ Estimated; speech symptoms first reported at this time.

Table S2. Cases and regions for qualitative identification of subregions with ubiquilin 2-only aggregates.

| Case type | Case | Region or sub-region | mtU2 aggs |
| --- | --- | --- | --- |
| **Hippocampus** |  |  |  |
| *UBQLN2*-linked ALS/FTD | MN17, V:7 | Dentate gyrus, Subiculum, Entorhinal cortex, Perirhinal cortex | **Yes** |
| Sporadic ALS | MN13, MN15 | Dentate gyrus, Subiculum, Entorhinal cortex, Perirhinal cortex |  |
| Non-neurodegenerative disease control | H238, H239, H245 | Dentate gyrus, Subiculum, Entorhinal cortex, Perirhinal cortex |  |
| **Spinal Cord** |  |  |  |
| *UBQLN2*-linked ALS/FTD | MN17 | Cervical segment 5, Thoracic segment 9, Lumbar segment 5 | **Yes** |
|  | V:7 | Cervical segment, Thoracic segment, Lumbar segment |  |
|  | IV:10 | Cervical segment |  |
| Sporadic ALS | MN12 | Cervical segment 8, Thoracic segment 9, Lumbar segment 5 |  |
|  | MN13 | Cervical segment 5, Thoracic segment 9, Lumbar segment 4 |  |
|  | MN15 | Cervical segment 5, Thoracic segment 9, Lumbar segment 2 |  |
| Non-neurodegenerative disease control | H211, H215, H248 | Cervical segment 1 |  |
| **Neocortex** |  |  |  |
| *UBQLN2*-linked ALS/FTD | MN17 | Motor cortex 0, Motor cortex 3, Sensory cortex 0, Sensory cortex 3, Superior frontal gyrus 1, Middle frontal gyrus 0, Middle frontal gyrus 2, Inferior frontal gyrus 1, Superior temporal gyrus 1, Middle temporal gyrus 0, Middle temporal gyrus 3, Inferior temporal gyrus 1, Superior parietal 1, Inferior parietal 1, Visual cortex 1, Visual cortex 2, Auditory cortex 1, Cingulate gyrus 0, Cingulate gyrus 4, Orbito-frontal gyrus 1 | **Yes** |
|  | V:7 | Motor cortex, Middle frontal gyrus |  |
|  | IV:10 | Cortical regions A-C^a^ |  |
| Sporadic ALS | MN13 | Motor cortex, Sensory cortex, Middle frontal gyrus 2, Middle temporal gyrus 2, Superior parietal 1, Visual cortex 2, Auditory cortex, Cingulate gyrus 0 |  |
|  | MN15 | Motor cortex 2, Sensory cortex 2, Middle frontal gyrus 2, Middle temporal gyrus 2, Superior parietal 1, Visual cortex 2, Auditory cortex, Cingulate gyrus 0, Cingulate gyrus 4 |  |
| Non-neurodegenerative disease control | H231, H238, H243 | Sensory-motor cortex 0 |  |
|  | H245 | Sensory-motor cortex 3 |  |
| **Striatum** |  |  |  |
| *UBQLN2*-linked ALS/FTD | MN17 | Caudate nucleus, Putamen, Ventral striatum | **Yes** |
|  | V:7 | Caudate nucleus, Putamen |  |
| Sporadic ALS | MN13, MN15 | Caudate nucleus, Putamen, Ventral striatum |  |
| Non-neurodegenerative disease control | H231, H238 | Caudate nucleus, Putamen, Ventral striatum |  |
| **Medulla** |  |  |  |
| *UBQLN2*-linked ALS/FTD | MN17 | Dorsal nucleus of vagus, Reticular formation (middle medulla), Raphe nuclei, Medial lemniscus, Inferior olivary nucleus, Medial accessory olivary nucleus, Medullary pyramids, Central grey, Gracile nucleus, Cuneate nucleus, Trigeminal nucleus, Reticular formation (lower medulla) | **Yes** |
|  | V:7 | Dorsal nucleus of vagus, Reticular formation, Raphe nuclei, Inferior olivary nucleus, Medial accessory olivary nucleus, Medullary pyramids |  |
| Sporadic ALS | MN13, MN15 | Dorsal nucleus of vagus, Reticular formation, Raphe nuclei, Medial lemniscus, Inferior olivary nucleus, Trigeminal nucleus, Medullary pyramids |  |
| **Substantia nigra** |  |  |  |
| *UBQLN2*-linked ALS/FTD | MN17  V:7  IV:10 | Substantia nigra 1  Substantia nigra  Midbrain-substantia nigra | **Yes** |
| Sporadic ALS | MN13  MN15 | Substantia nigra  Substantia nigra 1 |  |
| **Globus Pallidus** |  |  |  |
| *UBQLN2*-linked ALS/FTD | MN17  V:7 | Globus Pallidus  Globus Pallidus externa | **No** |
| Sporadic ALS | MN13, MN15 | Globus Pallidus |  |
| **Cerebellum** |  |  |  |
| *UBQLN2*-linked ALS/FTD | MN17 | Cerebellum 3 | **No** |
|  | V:7, IV:10 | Cerebellum |  |
| Sporadic ALS | MN13, MN15, | Cerebellum |  |
| Non-neurodegenerative disease control | H231, H243 | Cerebellum |  |

^a^Tissue was provided in blocks from unspecified cortical regions. Cortical region A was confirmed to be motor cortex.

Abbreviations. mtU2 aggs, ubiquilin 2-only aggregates present in *UBQLN2*-linked ALS/FTD cases in this region.

Table S3. Cases and regions for quantitative analysis of ubiquilin 2 and TDP-43 aggregates.

| Case type | Case | Region or sub-region |
| --- | --- | --- |
| **Hippocampus** |  |  |
| *UBQLN2*-linked ALS/FTD | MN17, V:7 | Dentate gyrus, Subiculum, Entorhinal cortex, Perirhinal cortex |
| Sporadic ALS | MN13, MN15, MN30 | Dentate gyrus, Subiculum, Entorhinal cortex, Perirhinal cortex |
| **Spinal Cord** |  |  |
| *UBQLN2*-linked ALS/FTD | MN17 | Cervical segment 5, Thoracic segment 9, Lumbar segment 5 |
|  | V:7 | Cervical segment, Thoracic segment, Lumbar segment |
|  | IV:10 | Cervical segment |
| Sporadic ALS | MN13 | Cervical segment 5, Thoracic segment 9, Lumbar segment 4 |
|  | MN15 | Cervical segment 5, Thoracic segment 9, Lumbar segment 2 |
|  | MN30 | Cervical segment 3, Thoracic segment 8, Lumbar segment 2 |
| **Motor cortex** |  |  |
| *UBQLN2*-linked ALS/FTD | MN17 | Motor cortex 0, Motor cortex 3 |
|  | V:7 | Motor cortex |
|  | IV:10 | Cortical region A^a^ |
| Sporadic ALS | MN13 | Motor cortex |
|  | MN15 | Motor cortex 2 |
|  | MN30 | Motor cortex |
| **Striatum** |  |  |
| *UBQLN2*-linked ALS/FTD | MN17 | Caudate nucleus, Putamen, Ventral striatum |
|  | V:7 | Caudate nucleus, Putamen |
| Sporadic ALS | MN13, MN15, MN30 | Caudate nucleus, Putamen, Ventral striatum |

^a^Tissue was provided in blocks from unspecified cortical regions. Cortical region A was confirmed to be motor cortex.

Table S4. Primary antibodies used for fluorescent immunohistochemistry.

| Primary antibody | Species (isotype) | Company | Catalogue # | Clonality | Dilution |
| --- | --- | --- | --- | --- | --- |
| **Qualitative analysis panels** | | | | | |
| *Spinal cord* | | | | | |
| Ubiquilin 2 | Mouse (Ig_G2a_) | Santa Cruz | SC-100612 | Monoclonal | 1:800 |
| pTDP-43 | Rat (IgG_2a_) | BioLegend | BL829901 | Monoclonal | 1:3,000 |
| SMI-32 (NFH) | Mouse (IgG_1_) | BioLegend | BL801701 | Monoclonal | 1:500 |
| MAP2 | Chicken (IgY) | Abcam | ab5392 | Polyclonal | 1:2,000 |
| *Histone H3 | Rabbit (IgG) | Abcam | ab1791 | Polyclonal | 1:500 |
| *All cortex* |  |  |  |  |  |
| Ubiquilin 2 | Mouse (IgG_2a_) | Santa Cruz | SC-100612 | Monoclonal | 1:800 |
| pTDP-43 | Rabbit (IgG) | CosmoBio | TIP-PTD-PO2 | Polyclonal | 1:4,000 |
| SMI-32 (NFH) | Mouse (IgG_1_) | BioLegend | BL801701 | Monoclonal | 1:500 |
| MAP2 | Chicken (IgY) | Abcam | ab5392 | Polyclonal | 1:2,000 |
| *Subcortical grey matter* | | | | | |
| Ubiquilin 2 | Mouse (IgG_2a_) | Santa Cruz | SC-100612 | Monoclonal | 1:800 |
| pTDP-43 | Rat (IgG_2a_) | BioLegend | BL829901 | Monoclonal | 1:3,000 |
| MAP2 | Chicken (IgY) | Abcam | ab5392 | Polyclonal | 1:2,000 |
| **Calbindin D-28k | Rabbit (IgG) | SWant | CB-38a | Polyclonal | 1:2,000 |
| **Leu-enkephalin | Rabbit (IgG) | Immunostar | 20066 | Polyclonal | 1:500 |
| *Histone H3 | Rabbit (IgG) | Abcam | ab1791 | Polyclonal | 1:500 |
| **Quantitative analysis panel** | | | | | |
| **Round 1** | | | | | |
| Ubiquilin 2 | Mouse (IgG_2a_) | Santa Cruz | SC-100612 | Monoclonal | 1:800 |
| Pan-TDP-43 | Rat (IgG_2a_) | BioLegend | BL808301 | Monoclonal | 1:300 |
| pTDP-43 | Rat (IgG_2a_) | BioLegend | BL829901 | Monoclonal | 1:3,000 |
| MAP2 | Chicken (IgY) | Abcam | ab5392 | Polyclonal | 1:2,000 |
| *Histone H3 | Rabbit (IgG) | Abcam | ab1791 | Polyclonal | 1:500 |
| **Round 2** |  |  |  |  |  |
| ***SMI-32 (NFH) | Mouse (IgG_1_) | BioLegend | BL801701 | Monoclonal | 1:500 |
| *Histone H3 | Rabbit (IgG) | Abcam | ab1791 | Polyclonal | 1:500 |
| **Confocal imaging panel** | | | | | |
| Ubiquilin 2 | Mouse (IgG_2a_) | Santa Cruz | SC-100612 | Monoclonal | 1:800 |
| Pan-TDP-43 | Rat (IgG_2a_) | BioLegend | BL808301 | Monoclonal | 1:300 |
| pTDP-43 | Rat (IgG_2a_) | BioLegend | BL829901 | Monoclonal | 1:3,000 |
| *Histone H3 | Rabbit (IgG) | Abcam | ab1791 | Polyclonal | 1:500 |
| ******Validation imaging panel** | | | | | |
| Ubiquilin 2 | Mouse (IgG_2a_) | Santa Cruz | SC-100612 | Monoclonal | 1:800 |
| Ubiquilin 2 | Mouse (IgG_1_) | Novus Biologicals | NBP2-25164 | Monoclonal | 1:800 |
| Pan-TDP-43 | Rat (IgG_2a_) | BioLegend | BL808301 | Monoclonal | 1:300 |
| pTDP-43 | Rat (IgG_2a_) | BioLegend | BL829901 | Monoclonal | 1:3,000 |
| *Histone H3 | Rabbit (IgG) | Abcam | ab1791 | Polyclonal | 1:500 |

*Histone H3 used only for case V:7 and omitted when calbindin or enkephalin used.

**Calbindin used for cerebellum and enkephalin used for globus pallidus.

***SMI-32 only included for spinal cord and motor cortex sections.

****Co-labelling of either Santa Cruz ubiquilin 2 + Novus ubiquilin 2; Santa Cruz ubiquilin 2 + pooled pan- and pTDP-43; or Novus ubiquilin 2 + pooled pan- and pTDP-43 was carried out for validation imaging.
Abbreviations. MAP2, microtubule-associated protein 2; pTDP-43, phospho (S409/410) TDP-43; NFH, neurofilament heavy chain.

Table S5. Secondary antibodies used for fluorescent immunohistochemistry.

| Secondary antibody or reagent | Conjugate | Company | Catalogue # | Dilution |
| --- | --- | --- | --- | --- |
| **Qualitative analysis panels** | | | | |
| *Spinal cord* | | | | |
| Goat anti-mouse IgG_2a_ | AlexaFluor® 488 | Invitrogen | A21131 | 1:500 |
| Goat anti-mouse IgG_1_ | AlexaFluor® 546 | Invitrogen | A21123 | 1:500 |
| Goat anti-chicken IgY | AlexaFluor® 594 | Invitrogen | A11042 | 1:500 |
| Goat anti-rat IgG | AlexaFluor® 647 | Jackson ImmunoResearch | 112.605.167 | 1:500 |
| *Goat anti-rabbit IgG | IRDye 800CW | LI-COR Biosciences | 92632211 | 1:500 |
| *All cortex* | | | | |
| Goat anti-mouse IgG_2a_ | AlexaFluor® 488 | Invitrogen | A21131 | 1:500 |
| Goat anti-mouse IgG_1_ | AlexaFluor® 546 | Invitrogen | A21123 | 1:500 |
| Goat anti-chicken IgY | AlexaFluor® 594 | Invitrogen | A11042 | 1:500 |
| Goat anti-rabbit IgG | IRDye 800CW | LI-COR Biosciences | 92632211 | 1:500 |
| *Subcortical grey matter* |  |  |  |  |
| Goat anti-mouse IgG | AlexaFluor® 488 | Jackson ImmunoResearch | 115-545-166 | 1:500 |
| Goat anti-chicken IgY | AlexaFluor® 594 | Invitrogen | A11042 | 1:500 |
| Goat anti-rat IgG | AlexaFluor® 647 | Jackson ImmunoResearch | 112.605.167 | 1:500 |
| Goat anti-rabbit IgG | IRDye 800CW | LI-COR Biosciences | 92632211 | 1:500 |
| **Quantitative analysis panel** | | | | |
| **Round 1** |  |  |  |  |
| *Goat anti-rabbit IgG | Biotin | Sigma-Aldrich | B7389 | 1:500 |
| *Streptavidin | AlexaFluor® 405 | ThermoFisher Scientific | S32351 | 1:200 |
| Goat anti-rat IgG | AlexaFluor® 488 | Jackson ImmunoResearch | 112-545-167 | 1:500 |
| Goat anti-mouse IgG | AlexaFluor® 594 | Jackson ImmunoResearch | 115-585-166 | 1:500 |
| Goat anti-chicken IgY | AlexaFluor® 647 | Invitrogen | A21449 | 1:500 |
| **Round 2** |  |  |  |  |
| *Goat anti-rabbit IgG | Biotin | Sigma-Aldrich | B7389 | 1:500 |
| *Streptavidin | AlexaFluor® 405 | ThermoFisher Scientific | S32351 | 1:200 |
| Goat anti-mouse IgG | AlexaFluor® 594 | Jackson ImmunoResearch | 115-585-166 | 1:500 |
| **Confocal imaging panel** |  |  |  |  |
| *Goat anti-rabbit IgG | Biotin | Sigma-Aldrich | B7389 | 1:500 |
| *Streptavidin | AlexaFluor® 405 | ThermoFisher Scientific | S32351 | 1:200 |
| Goat anti-rat IgG | AlexaFluor® 488 | Jackson ImmunoResearch | 112-545-167 | 1:500 |
| Goat anti-mouse IgG | AlexaFluor® 594 | Jackson ImmunoResearch | 115-585-166 | 1:500 |
| **Validation imaging panel** | | | | |
| *Co-label 1 (Santa Cruz ubiquilin 2 + Novus ubiquilin 2)* | | | | |
| Goat anti-mouse IgG_2a_ | AlexaFluor® 647 | Invitrogen | A21241 | 1:500 |
| Goat anti-mouse IgG_1_ | AlexaFluor® 488 | Invitrogen | A21121 | 1:500 |
| *Co-labels 2 and 3 (Santa Cruz ubiquilin 2 + pooled pan- and pTDP-43; Novus ubiquilin 2 + pooled pan- and pTDP-43)* | | | | |
| Goat anti-mouse IgG | AlexaFluor® 488 | Jackson ImmunoResearch | 115-545-166 | 1:500 |
| Goat anti-rat IgG | AlexaFluor® 647 | Jackson ImmunoResearch | 112-605-167 | 1:500 |
| *All sections* |  |  |  |  |
| *Goat anti-rabbit IgG | Biotin | Sigma-Aldrich | B7389 | 1:500 |
| *Streptavidin | AlexaFluor® 405 | ThermoFisher Scientific | S32351 | 1:200 |
| **All panels** |  |  |  |  |
| Hoechst 33342 | - | Cayman Chemicals | 15547 | 5 µg/mL |

* Goat anti-rabbit 800CW or goat anti-rabbit IgG biotin and streptavidin AlexaFluor® 405 used only for case V:7, with Hoechst 33342 being omitted.

### Supplementary Figures

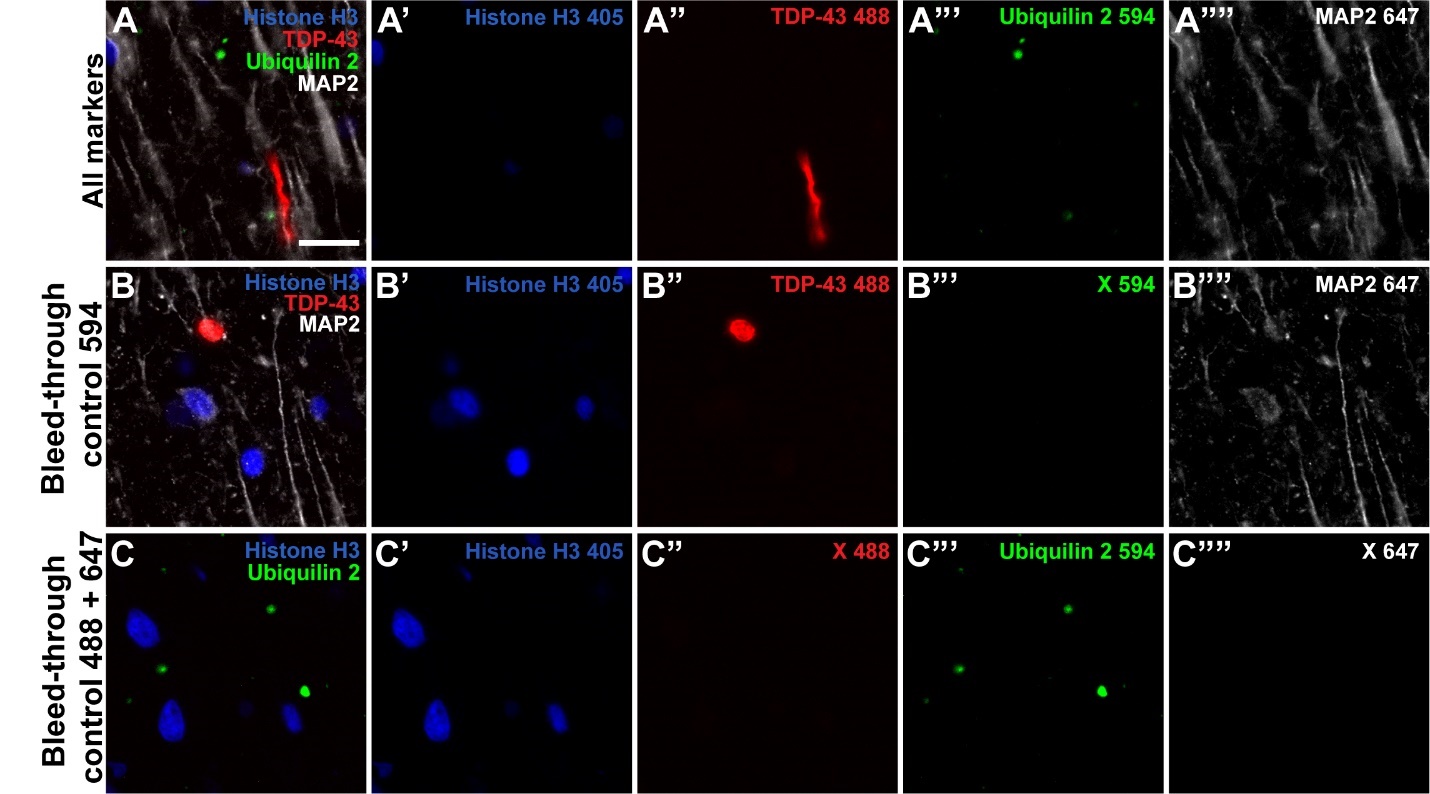

**Figure S1. Representative bleed-through control.**

Middle frontal gyrus sections from *UBQLN2* p.T487I-linked ALS/FTD case V:7 were labelled with all markers (A) or alternate markers (B and C) from Quantitative analysis panel Round 1 (See Tables S3 and S4). No bleed-through of fluorescence was observed from a given channel to the next channel that might be misidentified as co-localisation between ubiquilin 2, TDP-43, and/or MAP2. Scale bar 20 µm.

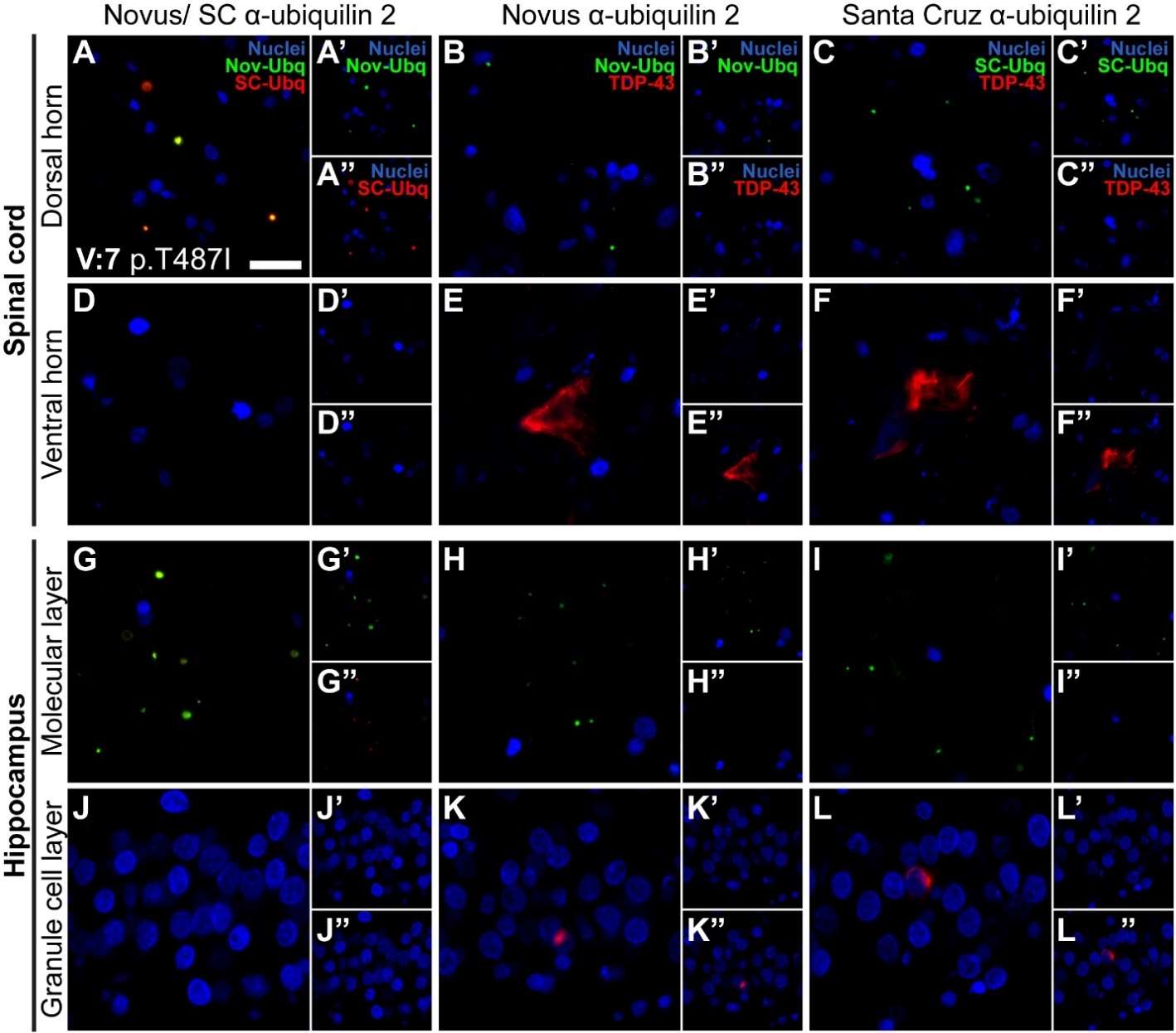

**Figure S2. Alternative ubiquilin 2 antibody labelling of *UBQLN2* p.T487I-linked ALS/FTD spinal cord and hippocampus.**

Validation of the monoclonal anti-ubiquilin 2 antibody used in this study (Santa Cruz #SC-100612, clone 5F5, IgG2a: SC-Ubq) was carried out by immunohistochemically co-labelling spinal cord (**A-F**) and hippocampus (**G-L**) sections from *UBQLN2* p.T487I-linked ALS/FTD case V:7 with another monoclonal anti-ubiquilin 2 antibody (Novus Biologicals #NBP2-25164, clone 6H9, IgG1: Nov-Ubq). To confirm the identified patterns of ubiquilin 2 and TDP-43 aggregation, we carried out three double-labels: Novus anti-ubiquilin 2 with Santa Cruz anti-ubiquilin 2; Novus anti-ubiquilin 2 with pooled anti-pan- and pTDP-43, and Santa Cruz anti-ubiquilin 2 with pooled anti-pan- and pTDP-43. Antibodies were visualised using AlexaFluor® 488 and 647 antibodies, respectively, to ensure spectral separation of markers. As seen with the Santa Cruz monoclonal anti-ubiquilin 2 antibody #SC-100612 in Figure 4, spinal cord sections from *UBQLN2*-linked cases demonstrated punctate aggregates in the dorsal horn that were positive for ubiquilin 2 only (**A-C**), with aggregates being immunoreactive for both the Santa Cruz and Novus anti-ubiquilin 2 antibodies. In the same sections, cytoplasmic aggregates in the ventral horn were positive for TDP-43 (**E”, F”,** red) and negative for ubiquilin 2 (**E’, F’,** green), irrespective of whether ubiquilin 2 was being visualised using the Santa Cruz or Novus antibody. The same pattern of ubiquilin 2 and TDP-43 aggregation was observed in the hippocampus, with ubiquilin 2-only aggregates identified in the molecular layer (**G-I**) and TDP-43-only aggregates identified in the granule cell layer (**J-L**), irrespective of whether ubiquilin 2 was visualised using the Santa Cruz or Novus antibody. Scale bars 20 µm.

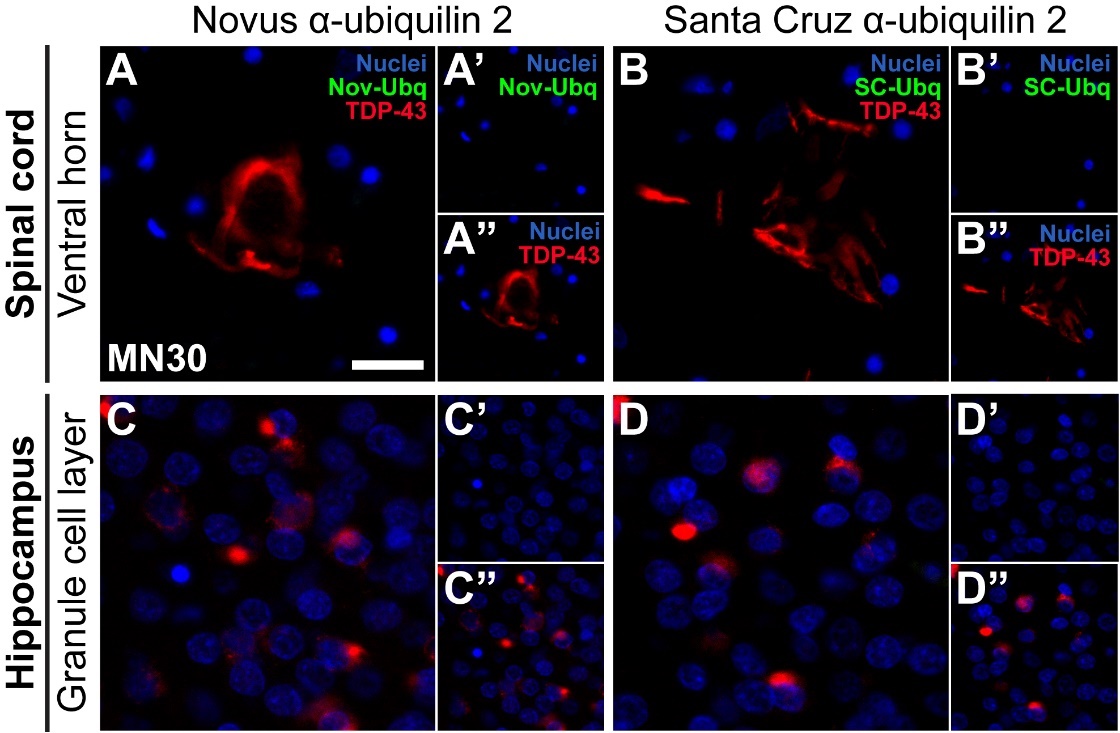

**Figure S3. Alternative ubiquilin 2 antibody staining of stage 4 sALS spinal cord and hippocampus.**

Validation of the monoclonal anti-ubiquilin 2 antibody used in this study (Santa Cruz #SC-100612, clone 5F5, IgG2a: SC-Ubq) was carried out by immunohistochemically co-labelling spinal cord (**A-B**) and hippocampus (**C-D**) sections from stage 4 sALS case MN30 with another monoclonal anti-ubiquilin 2 antibody (Novus Biologicals #NBP2-25164, clone 6H9, IgG1: Nov-Ubq). To confirm the identified patterns of ubiquilin 2 and TDP-43 aggregation, we carried out three double-labels: Novus anti-ubiquilin 2 with Santa Cruz anti-ubiquilin 2; Novus anti-ubiquilin 2 with pooled anti-pan- and pTDP-43, and Santa Cruz anti-ubiquilin 2 with pooled anti-pan- and pTDP-43. Antibodies were visualised using AlexaFluor® 488 and 647 antibodies, respectively, to ensure spectral separation of markers. Cytoplasmic aggregates in the ventral horn were positive for TDP-43 (**A”, B”,** red) and negative for ubiquilin 2 (**A’, B’,** green), irrespective of whether ubiquilin 2 was being visualised using the Santa Cruz or Novus antibody. The same pattern of ubiquilin 2 and TDP-43 aggregation was observed in the hippocampus, with TDP-43-only aggregates identified in the granule cell layer (**C-D**), irrespective of whether ubiquilin 2 was visualised using the Santa Cruz or Novus antibody. Scale bars 20 µm.

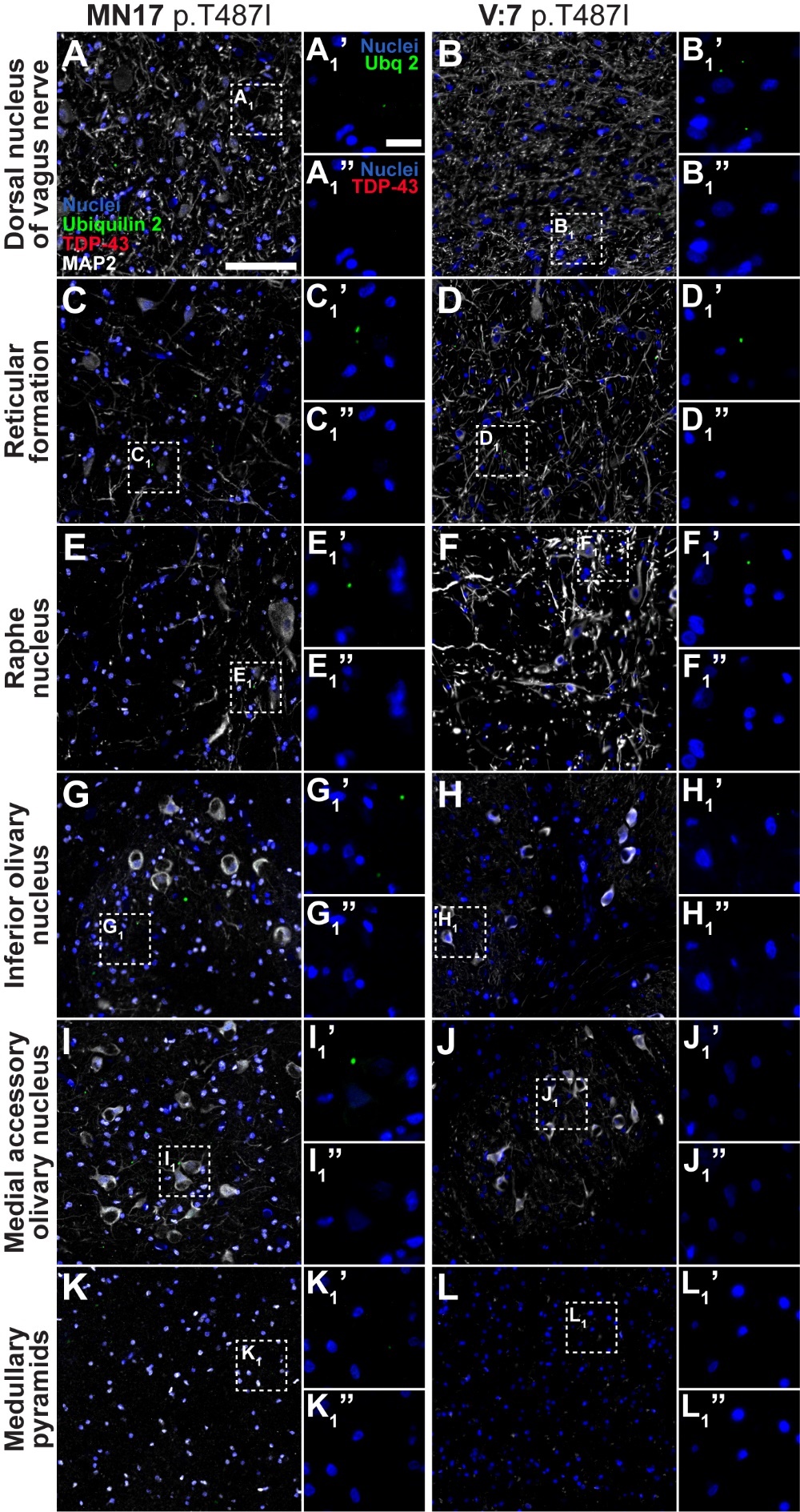

**Figure S4. Ubiquilin 2-only aggregates in the** ***UBQLN2* p.T487I-linked ALS/FTD middle medulla.**

Middle medulla sections from 2 *UBQLN2* p.T487I-linked ALS/FTD cases demonstrated punctate aggregates in the dorsal nucleus of the vagus nerve, reticular formation, raphe nucleus, and inferior olivary nucleus, that were positive for ubiquilin 2 only (**A_1_’-H_1_’,** green). In case MN17 but not case V:7, the medial accessory olivary nucleus and medullary pyramids also harboured punctate aggregates that were positive for ubiquilin 2 only (**I_1_’-L_1_’,** green). Scale bar in main images, 100 µm; in insets 20 µm.

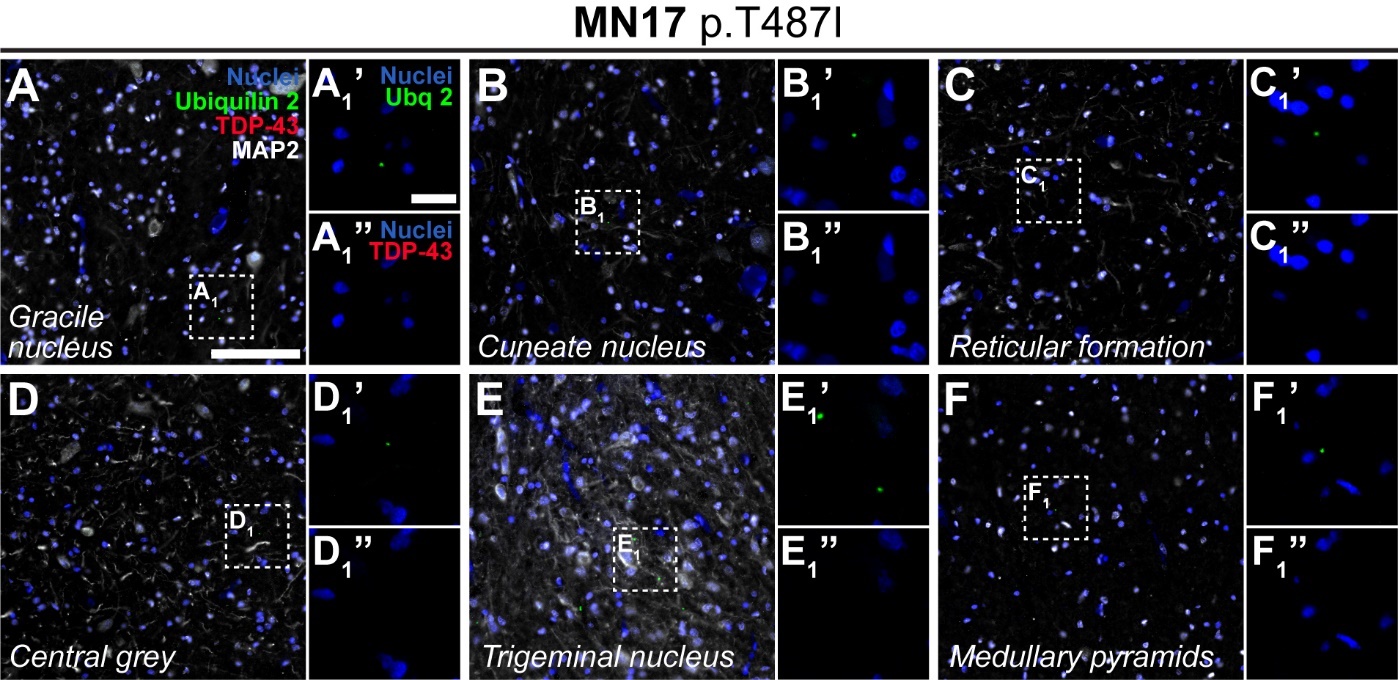

**Figure S5. Ubiquilin 2-only aggregates in the *UBQLN2* p.T487I-linked ALS/FTD lower medulla.**

Lower medulla sections from 1 *UBQLN2* p.T487I-linked ALS/FTD case demonstrated punctate aggregates in the gracile nucleus, cuneate nucleus, reticular formation, central grey, trigeminal nucelus, and medullary pyramids, that were positive for ubiquilin 2 only (**A’_1_-F’_1_,** green). Scale bar in main images, 100 µm; in insets 20 µm.

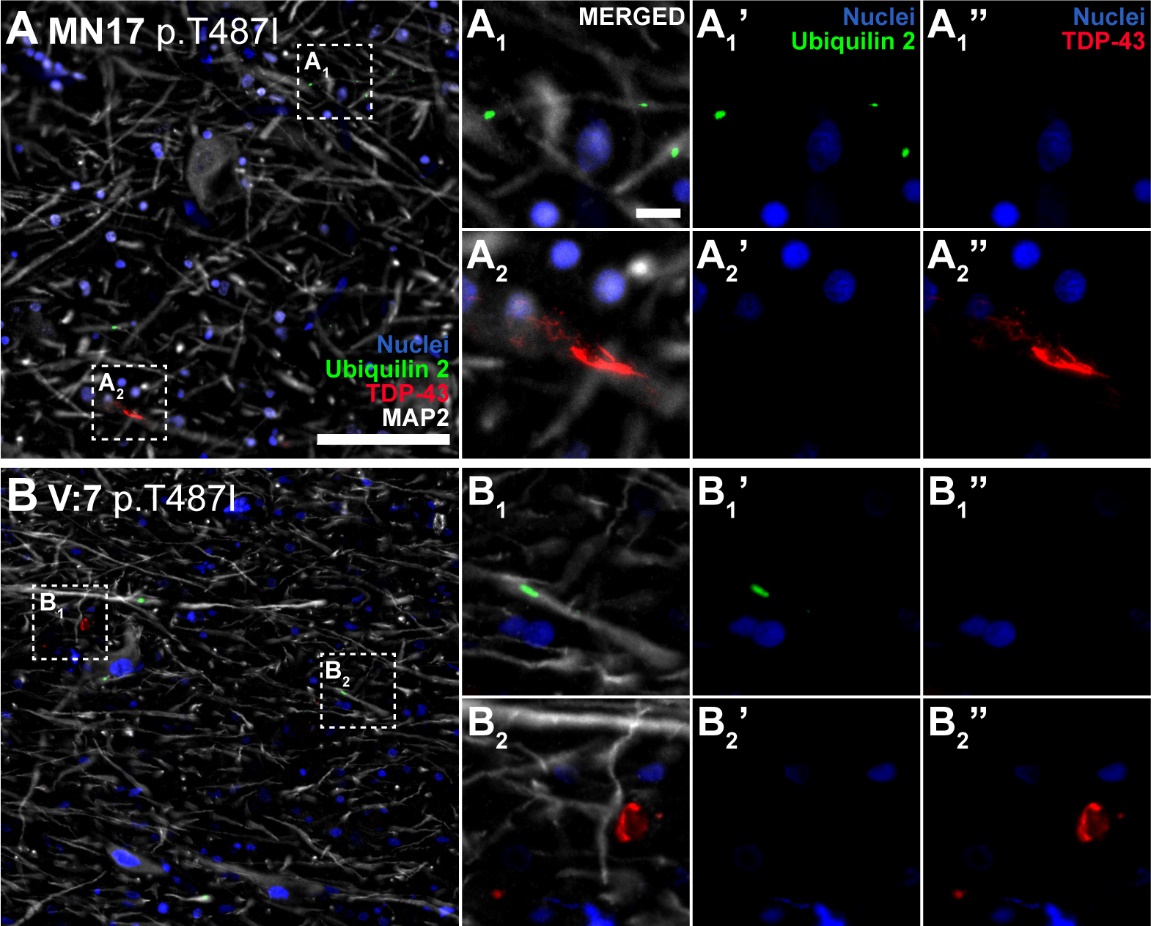

**Figure S6. Ubiquilin 2-only and TDP-43-only aggregates in the *UBQLN2* p.T487I-linked ALS/FTD substantia nigra.**

Substantia nigra sections from 2 *UBQLN2* p.T487I-linked ALS/FTD cases demonstrated punctate neuritic aggregates that were positive for ubiquilin 2 (**A_1_’-B_1_’,** green) or TDP-43 only (**A_2_”-B_2_”,** red), but not both. Scale bar in main images, 100 µm; in insets 10 µm.

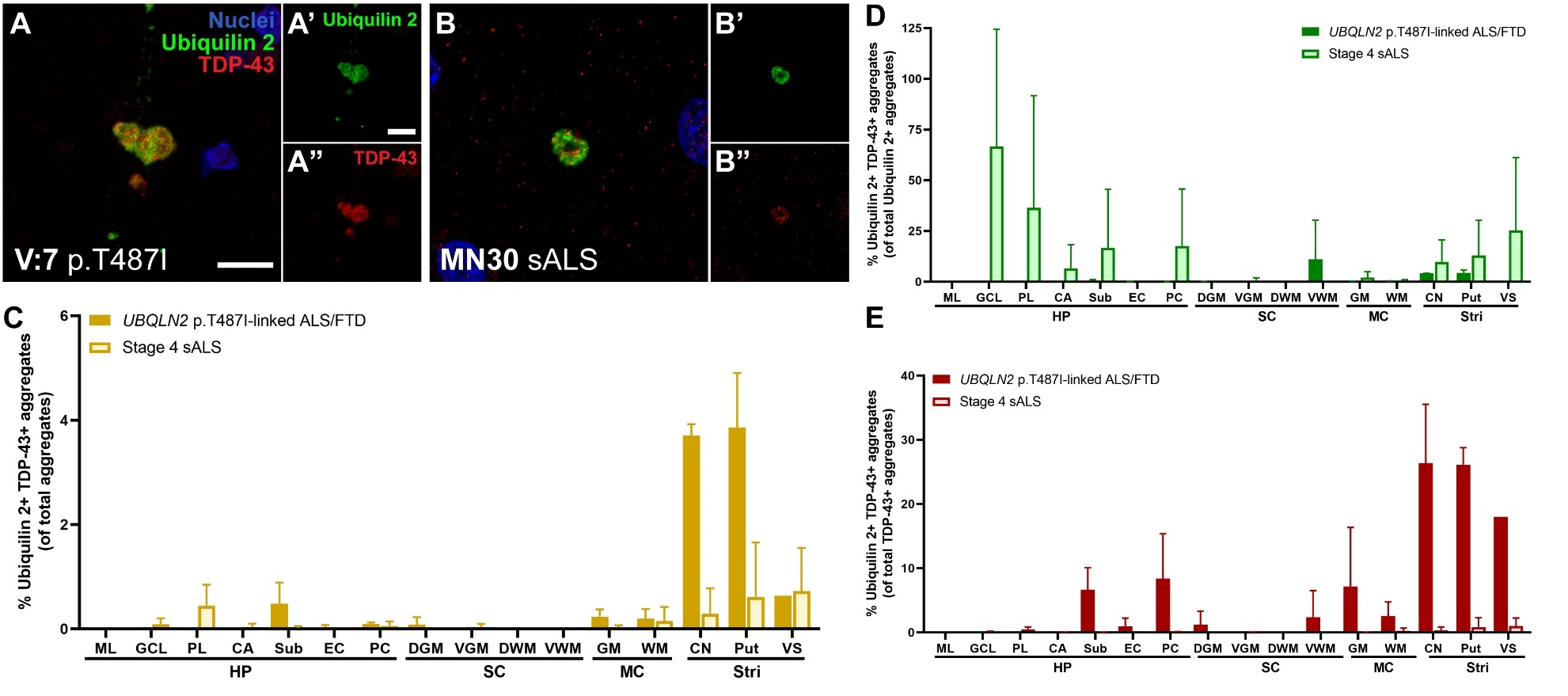

**Figure S7.** **Ubiquilin 2- and TDP-43-double immunopositive aggregates in the *UBQLN2* p.T487I-linked ALS/FTD and stage 4 sALS CNS.**

Confocal analysis of ubiquilin 2- and TDP-43-double immunopositive aggregates demonstrated that they were larger and more diffuse than punctate ubiquilin 2-only aggregates, similar in size to nuclei, and were present in both *UBQLN2* p.T487I-linked ALS/FTD and stage 4 sALS (**A,** **B**). Quantification of double immunopositive aggregates in sections from 2-3 *UBQLN2* p.T487I-linked ALS/FTD cases and 3 stage 4 sALS cases demonstrated these aggregates to be very rare as a proportion of all aggregates (<4%), and mostly found in striatum in *UBQLN2* p.T487I-linked ALS/FTD (**C**). Of all ubiquilin 2 aggregates, double immunopositivity with TDP-43 was common in stage 4 sALS hippocampus, but not in *UBQLN2* p.T487I-linked ALS/FTD (**D**). Of all TDP-43 aggregates, double immunopositivity with ubiquilin 2 was most common in *UBQLN2* p.T487I-linked ALS/FTD striatum (**E**). Scale bars 10 µm. Abbreviations: HP, hippocampus; ML, molecular layer; GCL, granule cell layer; PL, polymorphic layer; CA, cornu ammonis; Sub, subiculum; EC, entorhinal cortex; PC, piriform cortex; SC, spinal cord; DGM, dorsal grey matter; VGM, ventral grey matter; DWM, dorsal white matter; VWM, ventral white matter; MC, motor cortex; GM, grey matter; WM, white matter; Stri, striatum; CN, caudate nucleus; Put, putamen; VS, ventral striatum.

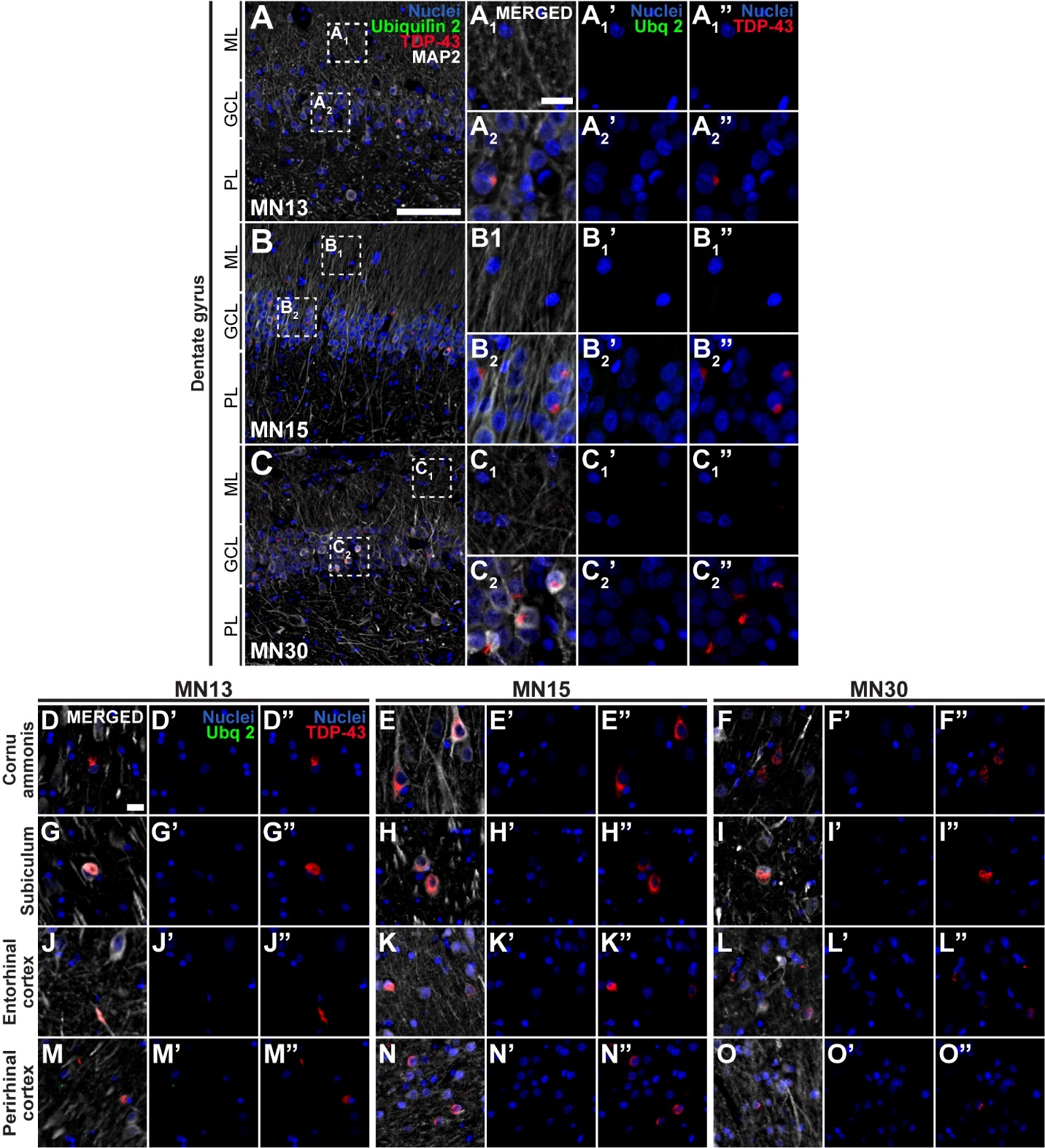

**Figure S8. Ubiquilin 2-only and TDP-43-only aggregates in the stage 4 sALS hippocampus.**

Hippocampal sections from 3 stage 4 sALS cases (**A-O”**) demonstrated cytoplasmic aggregates in the dentate gyrus and surrounding hippocampus that were positive for TDP-43 only (**A_1_”-O”,** red) and negative for ubiquilin 2 (**A_1_’-O’**). As shown in Figure 3L, quantification of the density of aggregates showed that ubiquilin 2-only aggregates were rare or absent but TDP-43-only aggregates were found throughout the hippocampal formation, particularly in the granule cell layer. Scale bars in A, B, C, 100 µm; others 20 µm. Abbreviations: ML, molecular layer; GCL, granule cell layer; PL, polymorphic layer.

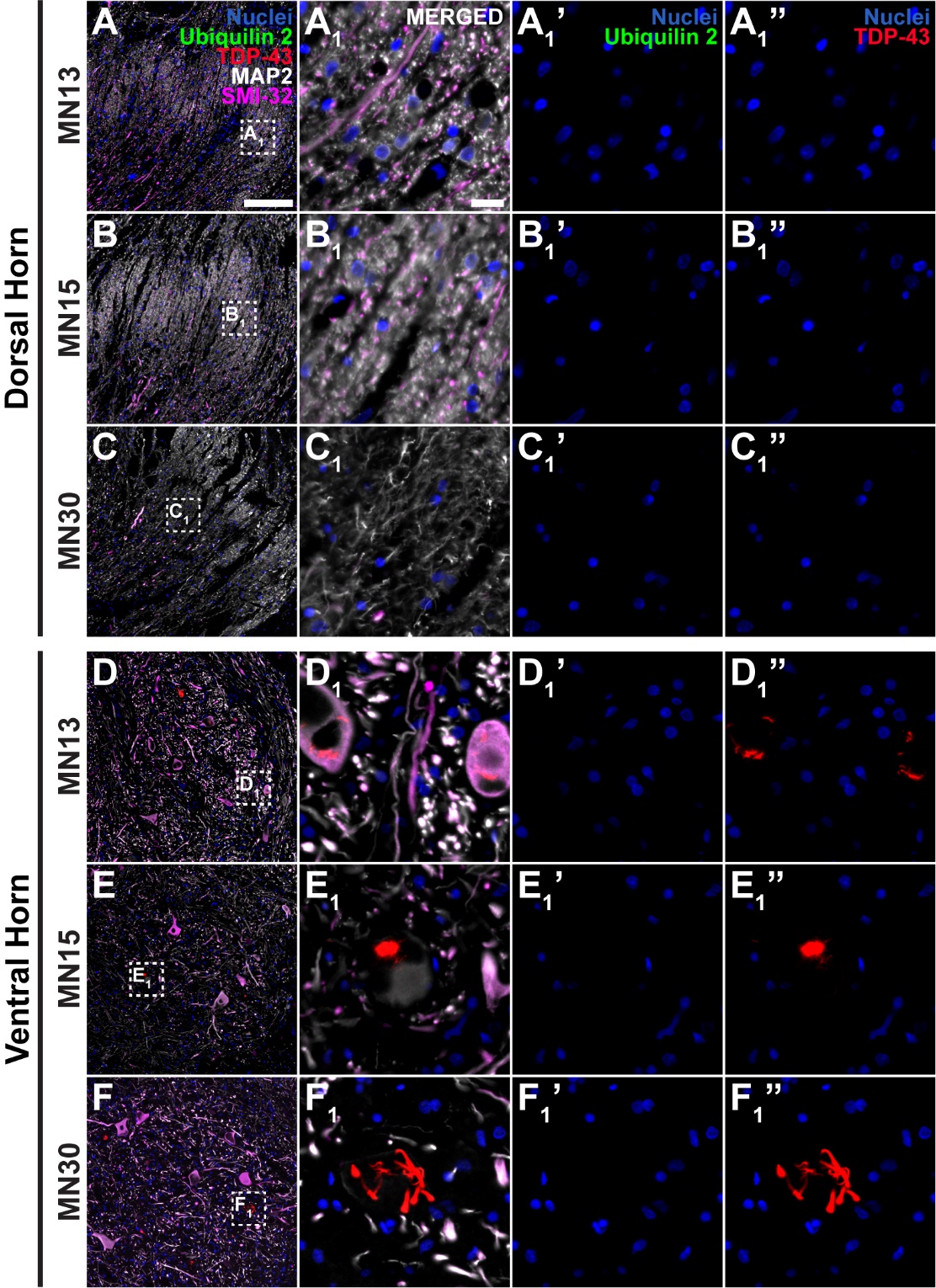

**Figure S9. Ubiquilin 2-only and TDP-43-only aggregates in the *UBQLN2* p.T487I-linked ALS/FTD spinal cord.**

Spinal cord sections from 3 stage 4 sALS cases (**A-F’_1_**) demonstrated cytoplasmic aggregates in the motor neurons of the ventral horn (magenta) that were positive for TDP-43 only (**D_1_”-F_1_”,** red) and negative for ubiquilin 2 (**D_1_’-F_1_’**, green). As shown in Figure 4H, quantification of the density of aggregates showed that ubiquilin 2-only aggregates were rare or absent, being most common in the dorsal grey matter, while TDP-43-only aggregates were abundant in the ventral grey matter. Scale bar in main images, 200 µm; in insets 20 µm.

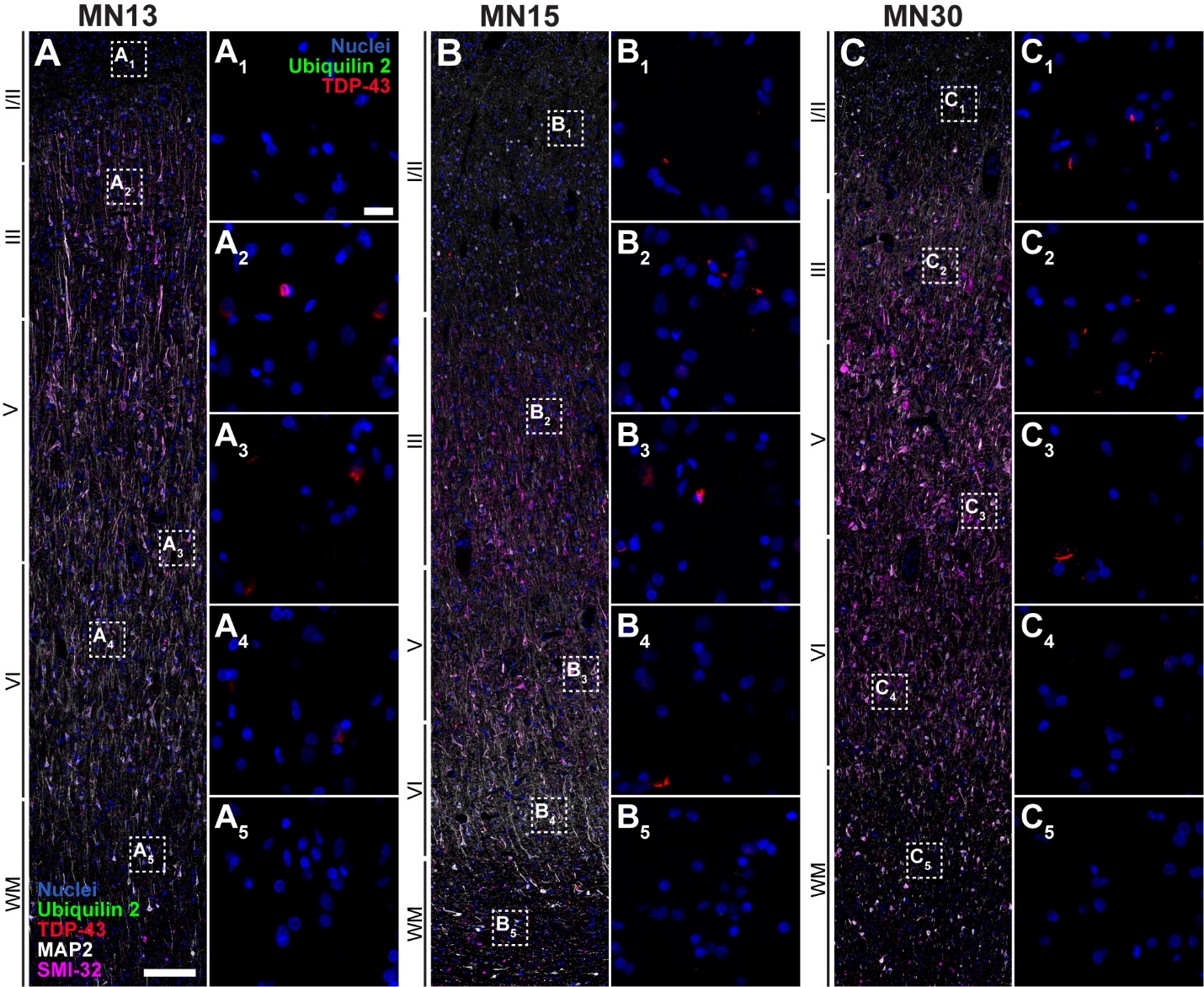

**Figure S10. Ubiquilin 2-only and TDP-43-only aggregates in the stage 4 sALS motor cortex.**

Motor cortex sections from 3 stage 4 sALS cases (**A-C_5_**) demonstrated cytoplasmic TDP-43-only aggregates (red) that were negative for ubiquilin 2. As shown in Figure 5E, quantification of the density of aggregates showed that ubiquilin 2-only aggregates (green) were very rare in (or absent from) all cortical layers, while TDP-43-only aggregates (red) were abundant throughout. Case MN30 showed occasional deposition of ubiquilin 2 in layers I-II and case MN15 showed greater deposition of TDP-43 in layers I-II. Scale bar in main images, 200 µm; in insets 20 µm. Abbreviations: WM, white matter.

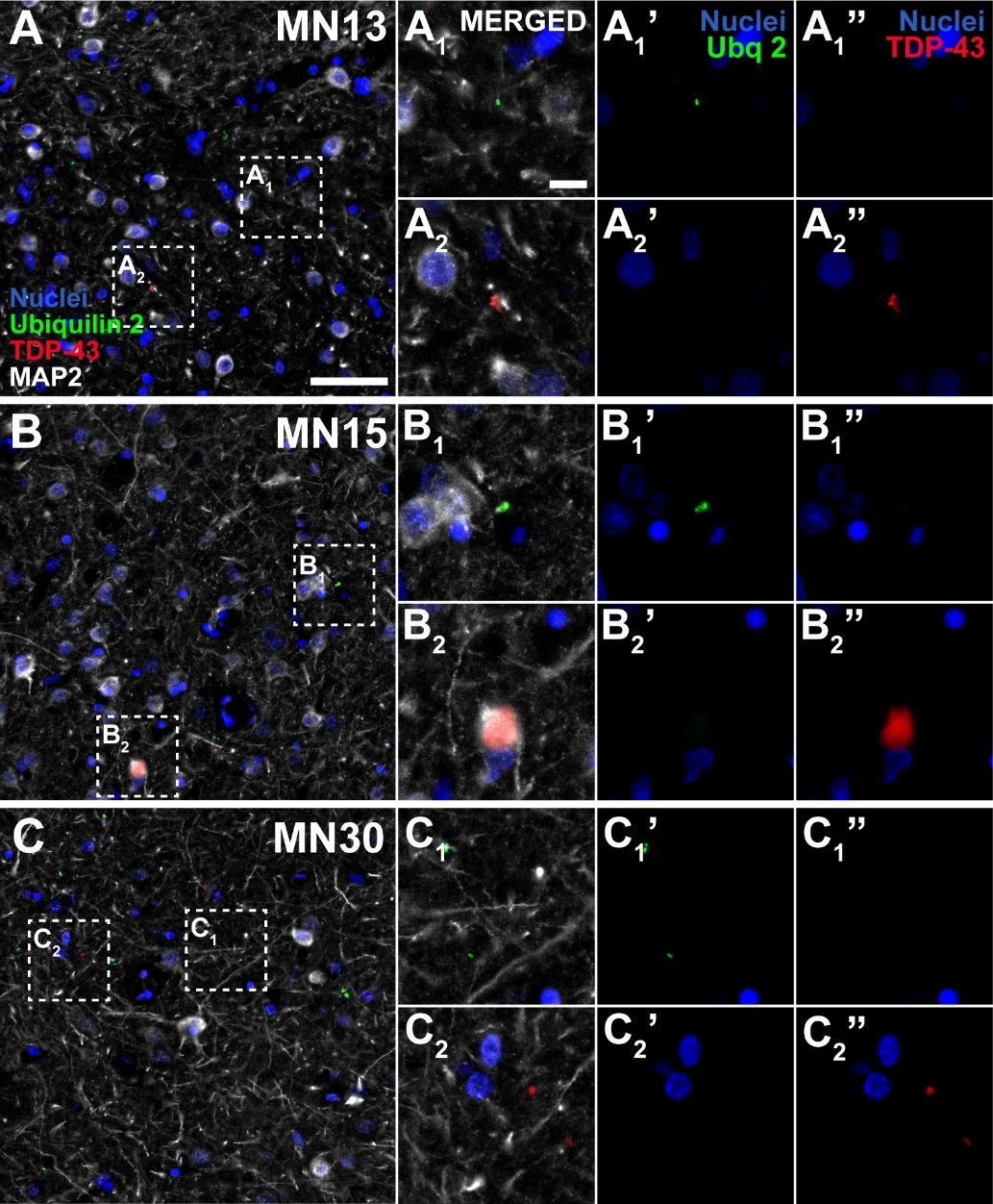

**Figure S11. Ubiquilin 2-only and TDP-43-only aggregates in the stage 4 sALS striatum.**

Striatal sections from 3 stage 4 sALS cases demonstrated punctate aggregates in the ventral striatum, that were positive for ubiquilin 2 only (**A’, A_1_’, A_2_’- C’, C_1_’, C_2_’**, green). Striatal sections also demonstrated cytoplasmic aggregates in that were positive for TDP-43 only (**A”, A_1_”, A_2_”- C”, C_1_”, C_2_”**, red). As shown in Figure 6G, quantification of the density of aggregates showed that ubiquilin 2-only aggregates (green) were most abundant in the putamen and ventral striatum of case MN30, while TDP-43-only aggregates (red) were abundant in these regions in case MN15. Scale bar in main images, 50 µm; in insets 5 µm.

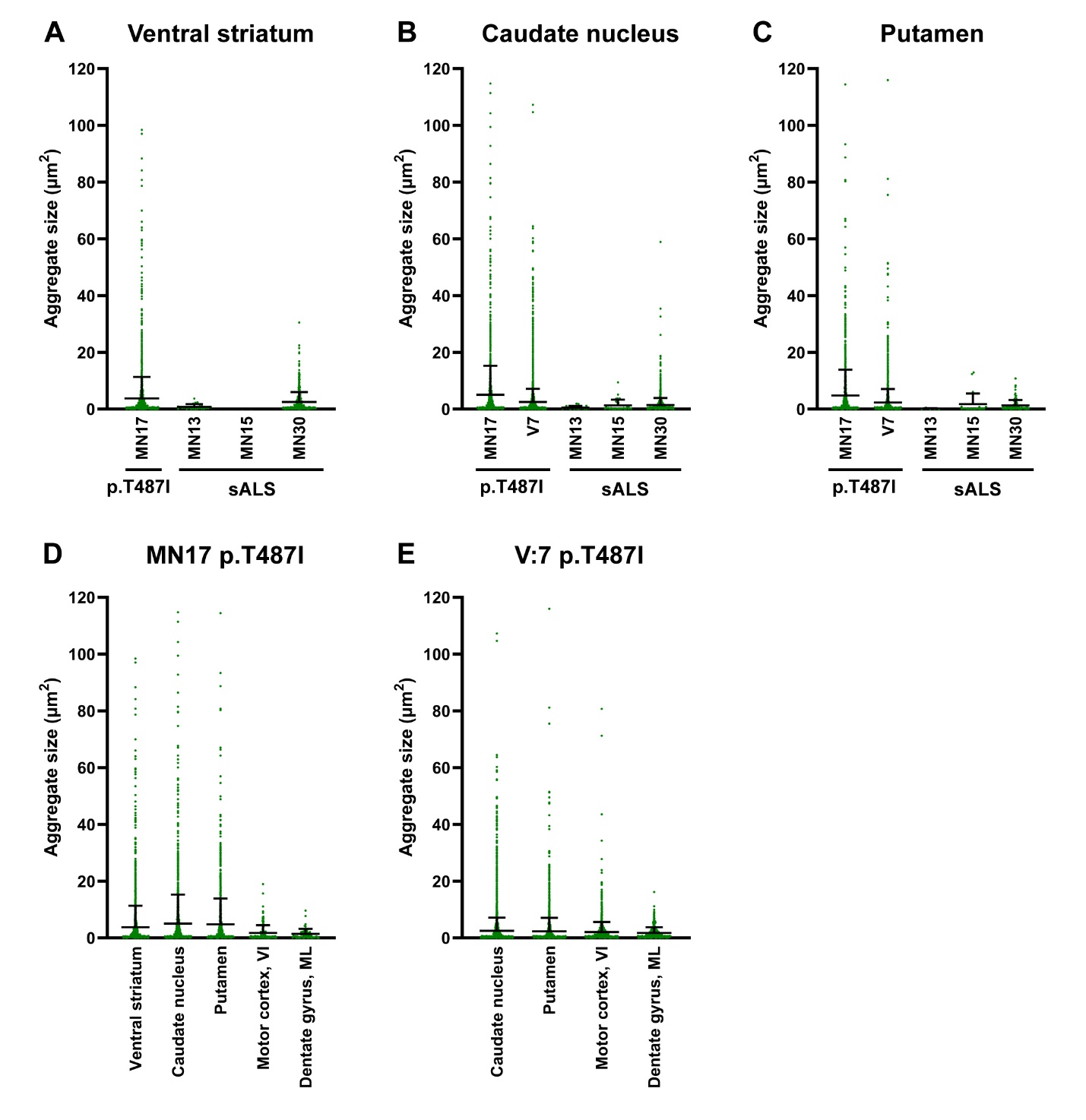

**Figure S12. Ubiquilin 2-only aggregate size distributions.**

Ubiquilin 2-only aggregate sizes were estimated in striatal sections from 2 *UBQLN2* p.T487I-linked ALS/FTD cases and 3 stage 4 sALS cases (**A-C**). Large ubiquilin 2-only aggregates (>40 µm^2^) were a feature of the striatum in *UBQLN2* p.T487I-linked cases (**D, E**).
